# Evaluating Best Practices for Isolating Pyrophilous Bacteria and Fungi from Burned Soil

**DOI:** 10.1101/2024.09.16.612975

**Authors:** Dylan J. Enright, Aishwarya Veerabahu, Ryan J. Quaal, Marely Vega, Anna Nguyen, Jenna Grindeland, James W.J. Randolph, Maria E. Ordonez, Sydney I. Glassman

## Abstract

A live microbial culture is invaluable to assess traits and functions, yet culturing immediately from fresh soil is logistically challenging and media selection is not trivial. Deeper ecological understanding of pyrophilous microbes and their traits is hampered by a lack of culture-based work causing researchers to rely heavily on community sequence analysis. To improve our understanding of isolating pyrophilous bacteria and fungi after wildfires from burned soil, we tested which: 1) soil storage method and 2) media retained highest genera richness and highest culturable viability as measured by CFUs retained. We tested four soil storage methods (dried, refrigerated at 4°C, stored at -80°C with or without glycerol) using a well-homogenized soil sample, plated microbes on rich media, and compared to fresh soil obtained 6 months after a severe California shrubland wildfire. From the fresh soils, we tested 3 media types: rich (Lysogeny Broth (LB) for bacteria; Malt Yeast Agar (MYA) for fungi), oligotrophic (Reasoner’s 2 Agar (R2A)) and media made from pyrogenic organic matter (PyOM). For bacteria, storing soil frozen at -80°C without glycerol yielded the highest genera richness and with glycerol preserved the most species. For fungi, storing soil at -80°C without glycerol preserved the most species but preserved equivalent richness of genera as fresh and 4°C soil. R2A yielded the highest bacterial and fungal genus richness but some species of interest were only captured with PyOM. Using a combination of these storage and media methods along with a few additional techniques (mushroom cultivation, smoke cultivation, etc.) from 2018-2022, we cultured >500 isolates (286 bacteria; 258 fungi) after 7 California wildfires for testing pyrophilous microbial traits.

**Importance:** Fires are increasing in frequency and severity across the globe, making the understanding of the traits of the pyrophilous microbes that survive and thrive post-fire and influence post-fire ecosystem regeneration critical and pressing. Yet, our ability to characterize the traits of pyrophilous microbes through genomics, transcriptomics, and phenotypic assays is hampered by lack of organisms in culture. Here, we debut a new medium created from pyrogenic organic matter that helps to cultivate fastidious pyrophilous microbes, and also describe a large-scale culture collection of pyrophilous bacteria and fungi (>500 isolates) that can be used for future collaborative research. We comprehensively test how soil storage and media type affect the cultivation of both bacteria and fungi, and for the first time, we test the best method to store soil to retain fungal diversity. Our work can be applied to improve culture collections of fastidious microbes from a variety of environments.

## Introduction

Although advances in the fields of molecular biology and DNA sequencing have freed microbiologists from the need to work with culturable species exclusively (1), few laboratory resources are as valuable as live cultures. Pure cultures are essential components for high quality whole-genome assembly (2, 3), genetic engineering experiments (4), and the phenotypic profiling of microorganisms (5–10). Additionally, microbial isolates greatly improve the interpretation of metabolomics data (11), and possession of a diverse culture collection can unlock a plethora of applied microbiological research directions (4, 12, 13). Once isolated and properly stocked, culture collections become a permanent resource for future microbiological research and generate new avenues for collaborative research (14–16). However, there are many hurdles to creating microbial culture collections and each collection has unique challenges (14).

The various factors that can be modified when attempting to culture microorganisms are as varied as there are microbes to isolate. For example, media selection may require grappling with an endless number of variables including consideration of the proper carbon sources, nitrogen and phosphorus content, salt concentrations, pH, and a host of other micronutrient considerations (13, 17, 18). One must also consider incubation temperature and duration, water and oxygen availability, and whether to alternate light and dark cycles or to make the media nutrient rich or poor (oligotrophic) (17–19). This complexity contributes to many microorganisms remaining uncultured (20). Though recent advances in our understanding of microbiology have begun to make the adage of “99% of microbes are unculturable” outdated (13, 18, 19, 21), culturing the majority of microorganisms still remains beyond our reach.

One group of understudied microorganisms that is of growing interest are the pyrophilous (or “fire-loving”) microbes. Pyrophilous microbes were rare or absent pre-fire but massively increase in abundance post-fire, sometimes persisting for many years post-fire (22–24), and influencing post-fire ecosystem recovery. While there are many studies identifying which microbes become more abundant post-fire, there are very few studies characterizing how pyrophilous microbes interact with their environment (25). Wildfires dramatically change landscapes and create legacy effects that can impact soil microorganisms and their overall ecosystems for many years (26–28). With rising global frequency and severity of wildfires across many ecosystems (29, 30), understanding how pyrophilous microbes modify and interact with their environments is critical. An important step to characterizing the ecology of pyrophilous microbes is laboratory testing of cultured isolates to provide data on how individual pyrophilous taxa compete for resources, cycle post-fire nutrients, and interact with the post-fire environment. However, there are numerous technological and logistical challenges to creating a culture collection of pyrophilous microbes.

Beyond the well-known hurdles of creating a microbial culture collection, attempting to culture microbes associated with wildfires presents a unique set of logistical challenges including proper soil storage. Wildfires are unpredictable natural phenomena. As such, it is often not possible to have the infrastructure for large-scale culturing efforts (personnel available, media selected and created, broth tubes or petri dishes ordered, culturing conditions decided and established) in place when a wildfire occurs, so post-fire soil samples may have to be stored for varying lengths of time prior to culturing. This is a logistical problem shared among many environmental microbial culturing efforts ranging from marine systems (31) to Antarctic soils (32). While many insights have been gained regarding these logistical hurdles such as culturing from samples as quickly as possible (14, 33), and methods for freezing samples that improve cultural viability, their efficacy is not always consistent and the viability of culturable microorganisms may be reduced depending on the soil storage method (33, 34).

Wildfires also radically alter environmental nutrient and resource landscapes (27, 35), making culture media selection challenging. Wildfire pyrolysis turns labile, easily accessible carbon sources into difficult to degrade pyrogenic organic matter (PyOM) (28, 36). PyOM is often enriched in polycyclic aromatic hydrocarbons (PAHs) that may require specialized metabolic pathways to process (12, 37–40). This can lead to accumulated char persisting in the environment for up to and exceeding 100 years (41). Additionally, wildfires deposit inorganic nitrogen that was previously locked in plant biomass through combustion and ash deposition (26, 35, 42). This high-nitrogen environment may alter the nutrient strategies that microorganisms employ post-fire and limit the microbes capable of surviving. For example, many species of mycorrhizal fungi are known to be nitrophobic (43), preventing them from establishing essential plant associations in high nitrogen environments. These nutrient shifts may result in commonly used laboratory media not providing appropriate resources to cultivate pyrophilous microbes.

Here, our goal was to create a large culture collection of pyrophilous microbes to use to test the traits that govern pyrophilous microbial adaptation to fire and the post-fire environment via biophysical assays and genomics. Keeping in mind the above described logistic and resource and nutrient-related challenges, we used soil from a local high severity California chaparral wildfire to test the best 1) soil storage methods and 2) laboratory media to retain the greatest richness and culturing viability of pyrophilous microbes. For soil storage, we tested culturing immediately from fresh soils versus soils stored for 3 months in four ways (air-dried, refrigerated at 4°C, and frozen at -80°C with and without glycerol). To reduce confounding variables, we plated all soil storage experiments on rich media. For media type, we tested culturing immediately from fresh soil onto rich (lysogeny broth (LB) agar for bacteria, malt-yeast agar (MYA) for fungi), oligotrophic (Reasoner’s 2 Agar (R2A)), or a media created in-house from PyOM. We tested the impact of our treatments on abundance (CFUs: Colony Forming Units) and microbial richness and composition (assessed via Sanger sequencing of 16S for bacteria or ITS for fungi; Figure 1). We hypothesized (H1) that other than fresh soil, air-dried soil would retain the greatest abundance and genera richness of pyrophilous bacteria and fungi since fires reduce soil moisture (35, 44), so we expected air drying soils would be the least dramatic disturbance for soil borne microorganisms. We further hypothesized (H2) that a combination of the rich and PyOM media would yield the greatest richness of pyrophilous microbes as the rich media could grow a wide range of bacteria and fungi and the PyOM media would capture those pyrophilous species with unique metabolic requirements specific to post-fire burned environments. Although we specifically sought to create a pyrophilous culture collection, the insights gleaned here may also aid environmental microbiologists in decision making about storage and media selection for many types of environmental culturing for fastidious microbes. Finally, the insights this study provided helped to guide our related work of creating a large and diverse collection of pyrophilous microorganisms of >540 isolates (286 bacteria; 258 fungi) from burned soils, mushrooms, and smoke derived from 7 California wildfires.

**Figure 1:**
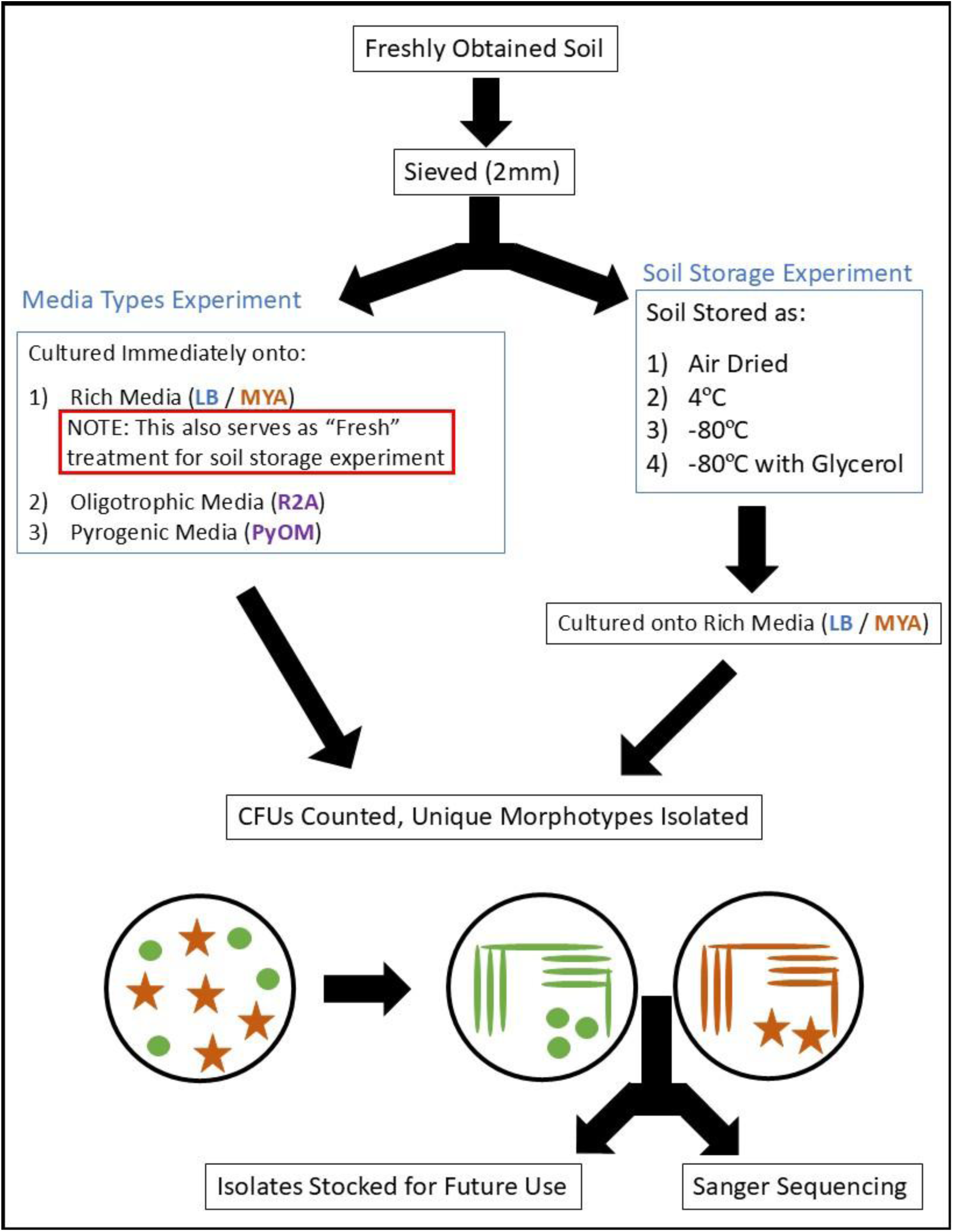
A conceptual diagram of the experimental design used in this experiment. “LB” is abbreviated for Lysogeny Broth, “MYA” is Malt Yeast Agar, “R2A” is Reasoner’s 2 Agar, and “PyOM” is Pyrogenic Organic Matter. Note that the cultures obtained from fresh soil plated onto rich media also serve as the “Fresh” storage condition for the soil storage experiment. Green circles and orange stars symbolize two different morphotypes collected from the same initial plate.

## Results

### Microbes Obtained

Throughout our soil storage and media selection experiments we isolated a visually diverse array of bacterial and fungal morphotypes (Figure 2). We cultured a total of 170 bacterial isolates representing 4 phyla, 10 orders, 31 genera and 83 taxa from burned soils collected 6 months after the 2020 El Dorado Fire. We also cultured 160 fungal isolates representing 3 phyla, 9 orders, 15 genera and 35 taxa (Table S1). Of these isolates, 138/170 bacterial isolates composed of 17 genera and 71 taxa and 152/160 fungal isolates composed of 8 genera and 30 taxa had literature support for inclusion as putative or confirmed pyrophilous taxa (22, 25, 27, 37, 38, 45–51) (Table S2). Putative pyrophilous taxa are those (or a close relative taxa) which appear repeatedly post-fire but have not been designated as pyrophilous in one or more publications yet. The putative pyrophilous taxa list includes more bacteria than fungi because the research on pyrophilous bacteria is more nascent than on pyrophilous fungi (24, 25, 45, 49), which dates back to 1909. We used the total (Table S1) and pyrophilous isolates (Table S2) to test the effect of media type and soil storage method on CFUs and microbial richness. In addition to testing storage methods and media selection, here we present our culture collection of 544 pyrophilous microbes (286 bacteria; 258 fungi) that can be used for testing of pyrophilous microbial traits, for example genomics, transcriptomics and phenotypic assays (52, 53), that is the result of our related work (from 2018-2022) in which we isolated bacteria and fungi from burned soils, mushrooms, and smoke of 7 California wildfires (Table S3). For bacteria, we had 286 isolates from 8 phyla, 16 orders, 35 genera, 76 taxa. For fungi, we had 258 isolates from 3 phyla, 11 orders, 13 genera, and 34 taxa. The full table includes the fires of origin, collection times, substrate cultures derived from, isolate names, FASTA sequences, and BLAST summary statistics (Table S3).

**Figure 2:**
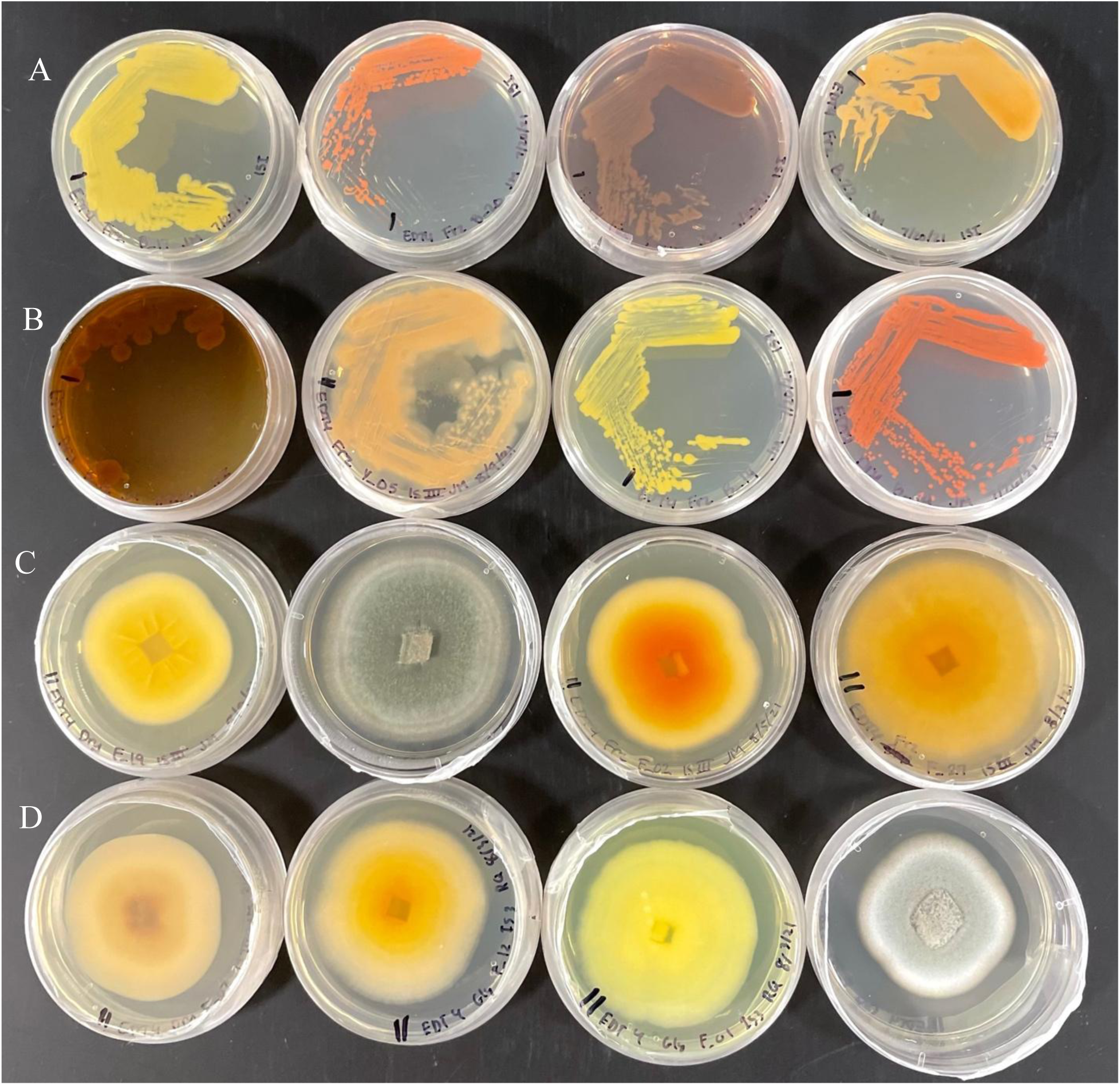
A snapshot of the morphological diversity of the cultures obtained from the El Dorado Fire across all soil storage and media type experiments. Identities were obtained via Sanger sequencing of the 16S or ITS regions for bacteria and fungi, respectively. Moving left to right across each row, the isolates shown are A)*Glutamicibacter bergerei, Arthrobacter agilis, Streptomyces mauvecolor, Kocuria rosea,* B) *Arthrobacter bussei, Holtermaniella festucosa, Pedobacter panaciterrae, Curtobacterium oceanosedimentum,* C) *Penicillium radiatolobatum, Penicillium fagi, Penicillium glabrum, Penicillium thomii,* D) *Penicillium chalabudae, Penicillium radiatolobatum, Penicillium murcianum, Penicillium adametzii*.

### Effect of Storage Method Selection on CFUs

Soil storage method significantly impacted the number of CFUs obtained per gram soil for both bacteria (ANOVA: F_1,4_ = 19.97, p < 0.001; Figure 3A) and fungi (ANOVA: F_1,4_ = 19.78, p < 0.001; Figure 3B). For both bacteria and fungi, soil stored air-dried (Bacteria: 2.31x10^5^ CFUs/gram, Fungi: 1.33x10^4^ CFUs/gram) and stored at - 80°C with glycerol (Bacteria: 1.31x10^5^ CFUs/gram, Fungi: 6.69x10^3^ CFUs/gram) retained the fewest average CFUs/gram soil. For bacteria, soils stored at 4°C (6.98x10^5^ CFUs/gram) or at - 80°C without glycerol (6.39x10^5^ CFUs/gram) were not significantly different from fresh soils (5.73x10^5^ CFUs). For fungi, fresh soils had the highest average CFUs/gram soil (3.07x10^4^ CFUs/gram), followed by soil frozen at -80°C without glycerol (2.66x10^4^ CFUs/gram), and then stored at 4°C (2.16x10^4^ CFUs/gram).

**Figure 3:**
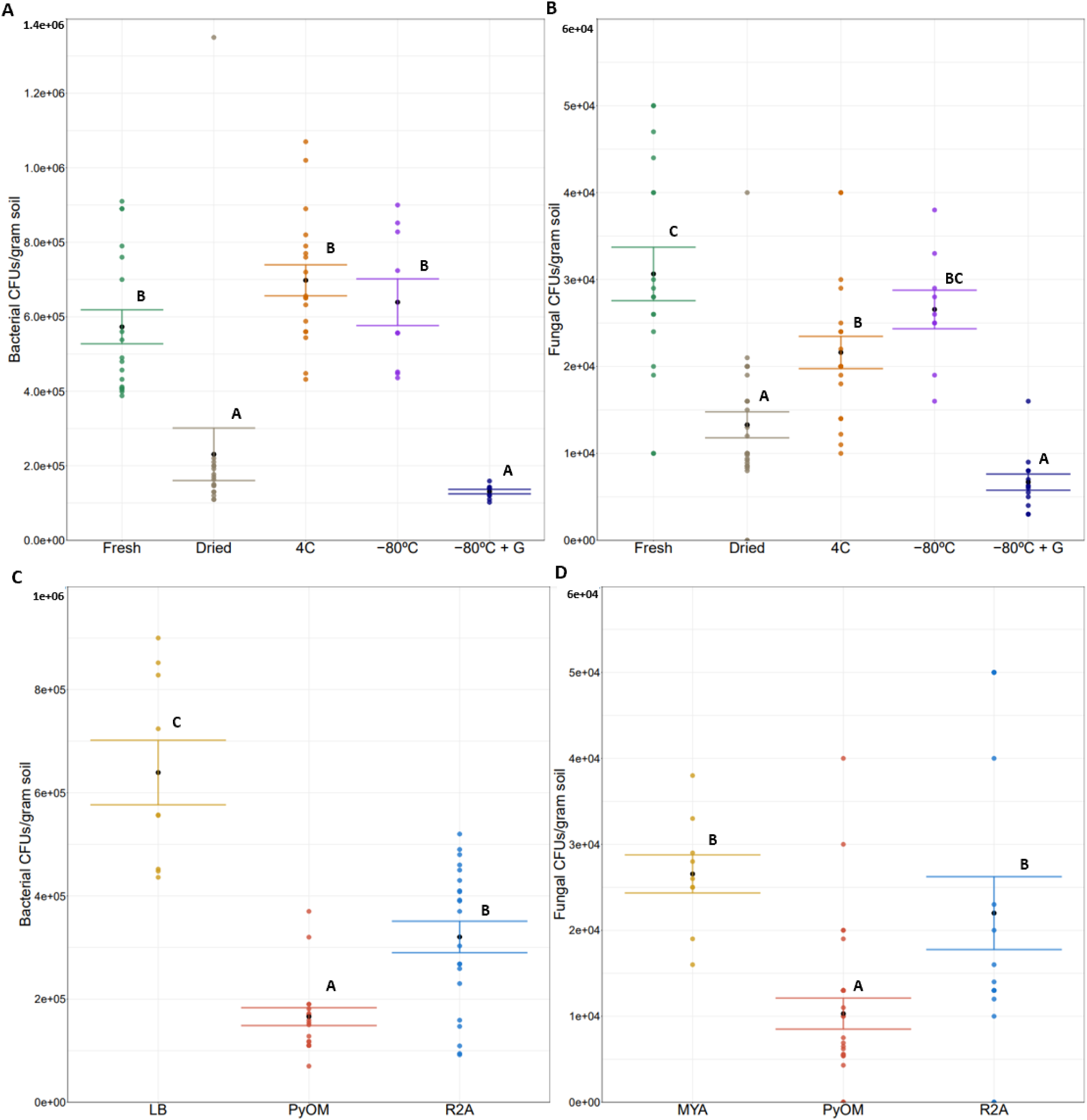
Calculated capturable CFUs per gram soil for A) bacteria and B) fungi among the tested soil storage methods and C) bacteria and D) fungi across tested media types. Points and bars are colored by treatment with bars indicating standard error of themean and black points indicating the mean CFUs per gram soil for each treatment. Significance was tested using ANOVA and Tukey Post-Hoc lettering has been applied. LB = Lysogeny Broth; PyOM = Pyrogenic Organic Matter; R2A = Reasoner’s 2 Agar, and G stands for glycerol.

### Effect of Media Selection on CFUs

Media type significantly impacted the number of CFUs obtained per gram of fresh soil for both bacteria (ANOVA: F_1,2_ = 39.43, p < 0.001; Figure 3C) and fungi (F_1,2_ = 7.77, p < 0.01; Figure 3D). For both bacteria and fungi, PyOM media yielded the fewest average CFUs per gram (Bacteria: 1.66x10^5^; Fungi: 1.03x10^4^), and rich media (LB for bacteria and MYA for fungi) had the highest average CFUs per gram (Bacteria: 5.73x10^5^; Fungi: 3.07x10^4^). R2A media had the second highest per gram average CFU counts for bacteria (3.20x10^5^) and for fungi, CFU counts (2.20x10^4^) were statistically equivalent to MYA.

### Effect of Storage Method Selection on Bacterial Diversity

Soil storage method strongly affected richness of bacterial genera with fresh soil retaining the highest total (Figure 4A) and frozen soil retaining the highest pyrophilous richness (Figure 4B). Dried soil retained the lowest richness of bacterial genera among both total (Figure 4A) and pyrophilous isolates (Figure 4B). Across all soil storage treatments, we cultured a total of 21 bacterial genera (Figure S2) and 64 taxa (Figure S3), and we found literature support to classify 14 genera (Figure 6A) and 51 taxa (Figure S4) as pyrophilous or putatively pyrophilous. Considering total bacterial taxa, fresh and frozen soil were equally rich (62% of genera), followed by soil frozen with glycerol (57% of genera), 4°C soil (48% of genera) with dried soil as least rich (38% of genera). Only 3 genera (*Arthrobacter, Curtobacterium, Streptomyces*) were isolated across all 5 storage treatments whereas 8 genera (*Brevibacterium, Fontibacillus, Glutamicibacter, Massilia, Ornithinimicrobium, Paeniglutamicibacter,* an unidentified Firmicute, and *Pseudoarthobacter*) were isolated from a single treatment (Figure S2). Finally, 64% of taxa were isolated one time with only one taxon best identified as *Curtobacterium oceanosedimentum* observed in all storage treatments (Figure S3). Among the pyrophilous subset (Figure 4B, S4), frozen soil was the most rich (85% of genera), followed by fresh soil (62% of genera), soil frozen with glycerol (54% of genera), 4°C soil (54% of genera) with dried soil as least rich (38% of genera).

**Figure 4:**
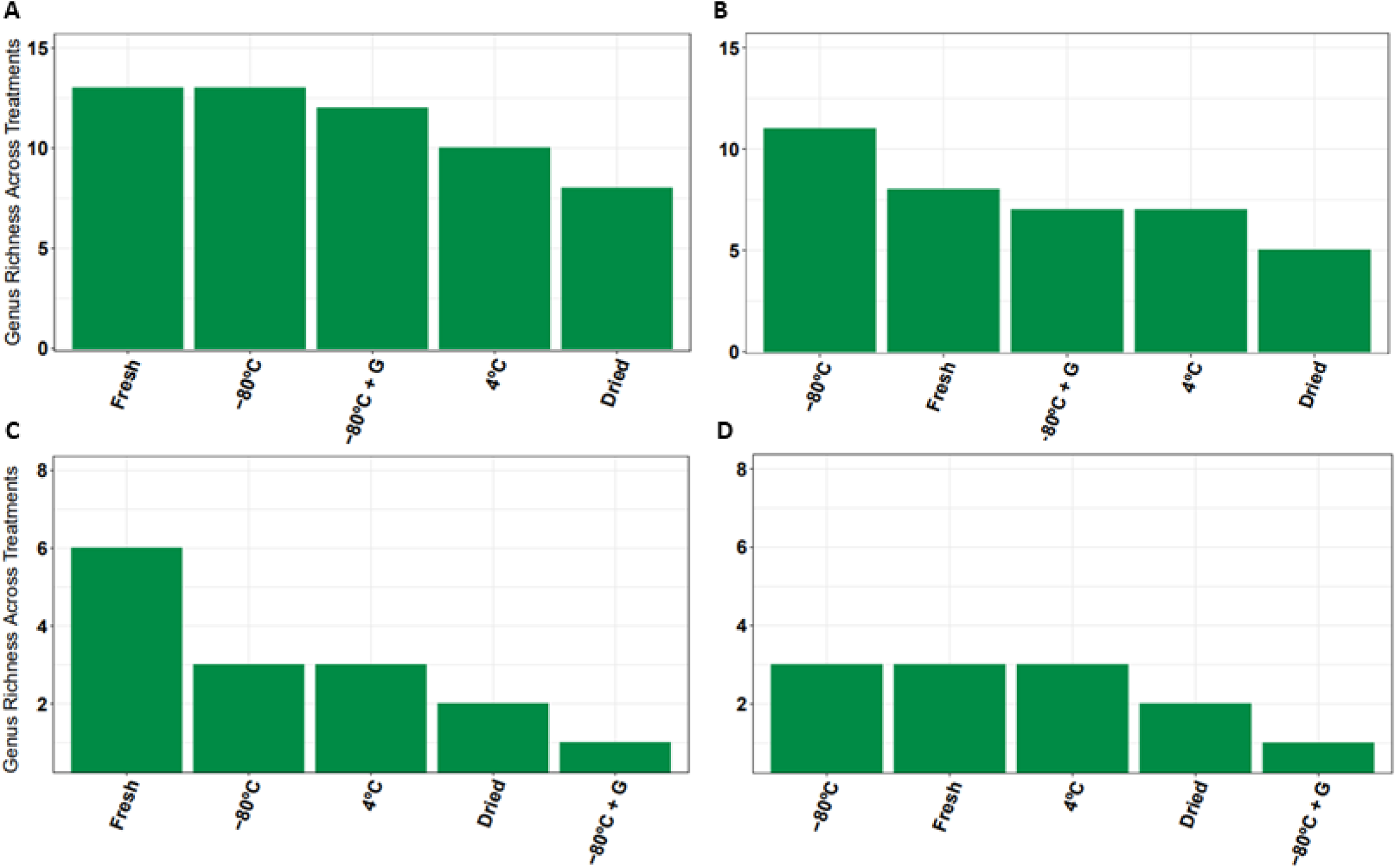
Summarized counts of A) total bacterial, B) pyrophilous bacterial, C) total fungal, and D) pyrophilous fungal genera across four soil storage methods compared to fresh soil when isolated on LB for bacteria or MYA for fungi. Treatments are ordered by highest amount of genera. G = glycerol.

### Effect of Storage Method Selection on Fungal Diversity

Soil storage method strongly affected fungal richness with fresh soil retaining the highest total genera (Figure 4C) and fresh, frozen and refrigerated retaining equal numbers of pyrophilous genera (Figure 4D). Soil frozen with glycerol retained the lowest richness of fungal genera among total (Figure 4C) and pyrophilous isolates (Figure 4D). Across all storage treatments, we cultured a total of 6 fungal genera (Figure S5) and 23 fungal taxa (Figure S6), of which we found literature support as pyrophilous for 3 genera (Figure 6B) and 20 taxa (Figure S7). For total fungi, fresh soil was the most rich, capturing 100% of all 6 genera (*Aspergillus, Holtermaniella, Ochrocladiosporium*, *Penicillium*, and 2 uncultured Helotiales), followed by frozen and refrigerated soil capturing 50% of the genera, and 30% in dried soil (Figure S5). Soil frozen with glycerol performed the worst, capturing only *Penicillium* (17% of the total; Figure S5). For pyrophilous fungi, fresh, frozen, and refrigerated soil all captured 100% of the 3 genera (*Aspergillus, Holtermanniella, Penicillium*), whereas dried captured 2 (*Aspergillus, Penicillium*) and frozen with glycerol only captured 1 (*Pencillium*; Figure 6B).

### Effect of Media Selection on Bacterial Diversity

Media type strongly affected bacterial richness with R2A and LB both capturing the highest numbers of total bacterial genera (Figure 5A), R2A yielding the highest richness of pyrophilous genera (Figure 5B), and PyOM capturing the fewest total and pyrophilous bacterial genera. Across all media types from fresh soil, we cultured 24 total bacterial genera (Figure S8), 61 total bacterial taxa (Figure S9), and 16 pyrophilous bacterial genera (Figure 7A) and 49 pyrophilous bacterial taxa (Figure S10). Among total bacteria, LB and R2A obtained equally rich communities (58% of genera), with the least in PyOM (33% of genera). Among pyrophilous bacteria, R2A captured the greatest richness (69% of genera), followed by LB (56% of genera) with PyOM yielding the lowest richness (44% of genera). Although PyOM media yielded the lowest diversity, it was the only media to isolate some important pyrophilous bacterial genera like *Noviherbaspirillum* and *Peribacillus* (46, 47, 49, 52) and interestingly, R2A was the only media to capture *Massilia* (Figure 7A) suggesting a more oligotrophic lifestyle than previously thought (45, 49).

**Figure 5:**
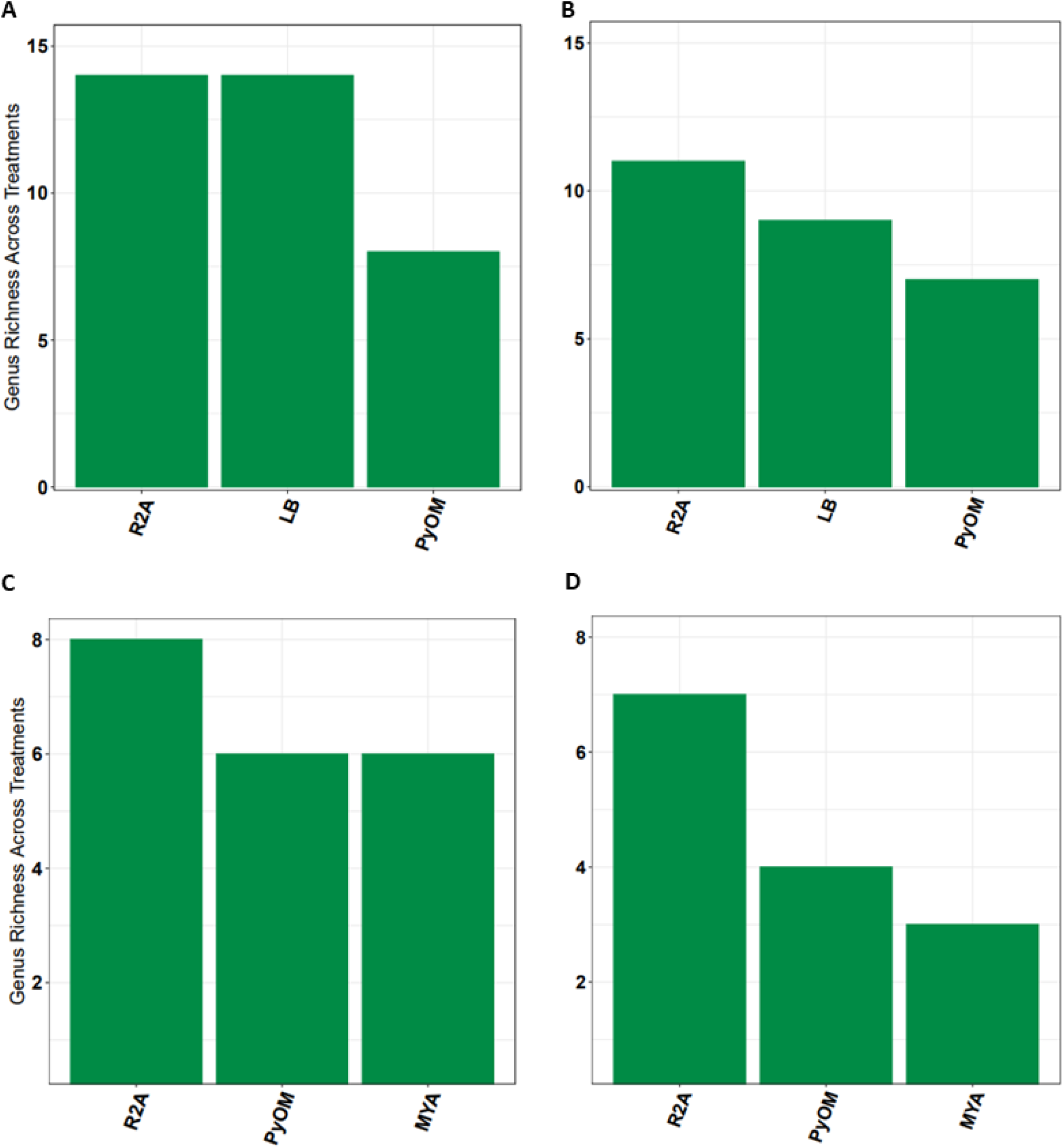
Summarized counts of A) total bacterial, B) pyrophilous bacterial, C) total fungal, and D) pyrophilous fungal genera (green) obtained from fresh soil tested across three media types. Treatments are ordered by highest number of genera. LB = Lysogeny Broth, MYA = Malt Yeast Agar, R2A = Reasoner’s 2 Agar, PyOM = Pyrogenic Organic Matter.

**Figure 6:**
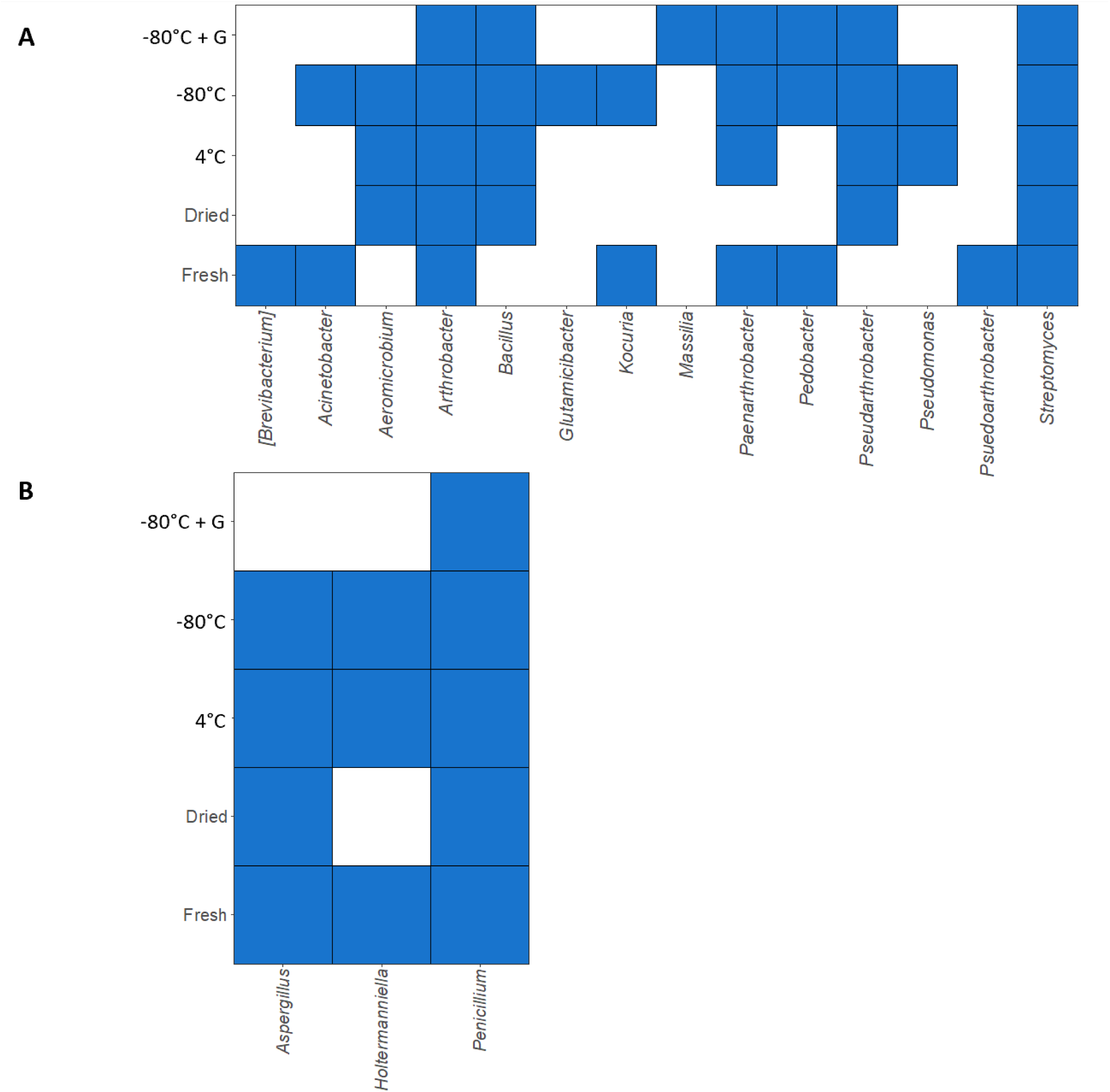
Heatmap-style visualization showing the diversity of pyrophilous A) bacterial and B) fungal genera across the soil storage experiment. A filled blue square indicates the presence of the genus on the x axis in the soil storage method indicated on the y axis. G = glycerol. Genera are ordered in alphabetical order, other than [Brevibacterium] which is in brackets due to controversy over its taxonomy.

**Figure 7:**
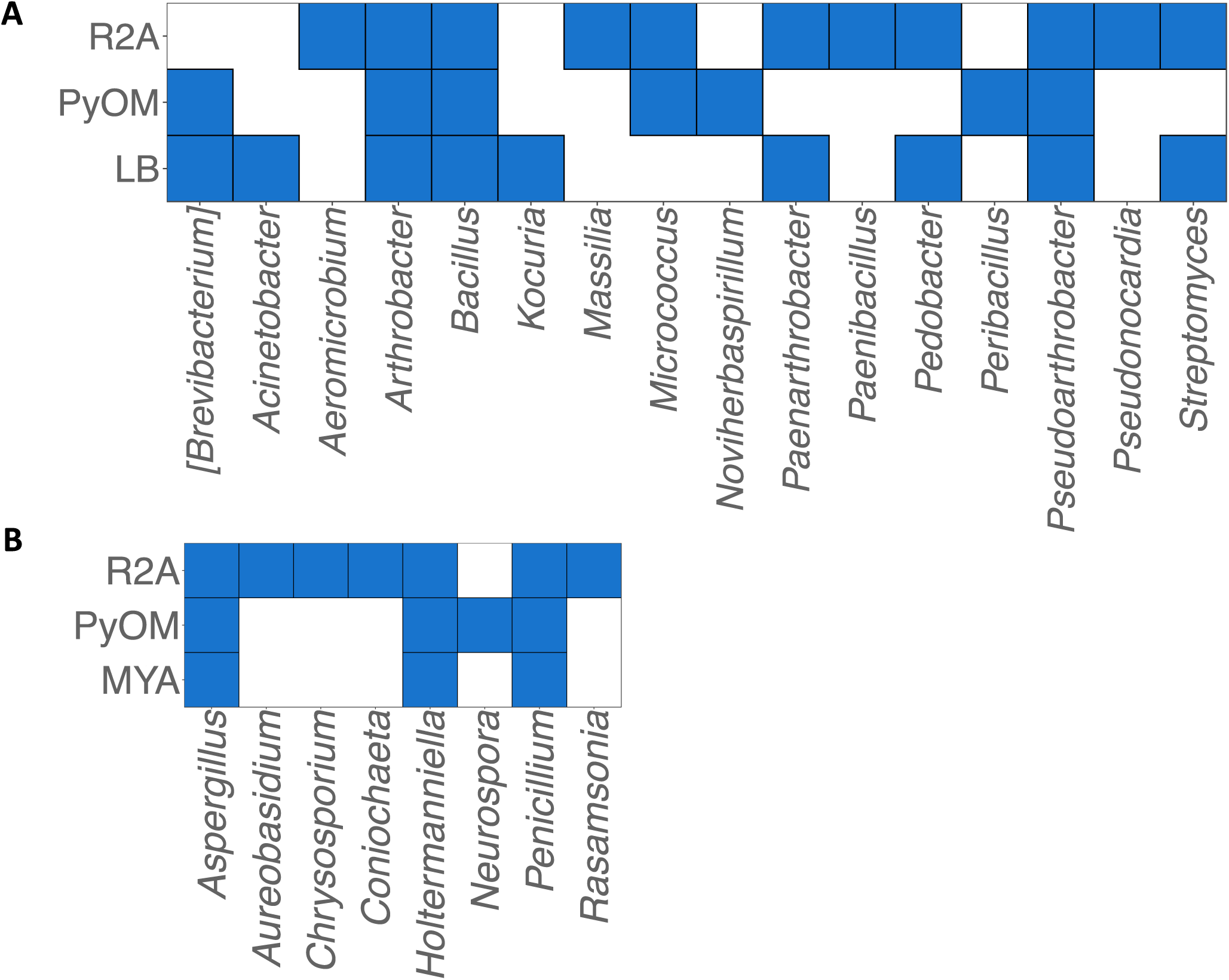
Heatmap-style visualization showing the diversity of pyrophilous A) bacterial and B) fungal genera across the media types experiment. A filled blue square indicates the presence of the genus on the x axis in the media indicated on the y axis. LB = Lysogeny Broth, MYA = Malt Yeast Agar, R2A = Reasoner’s 2 Agar, PyOM = Pyrogenic Organic Matter. Genera are ordered in alphabetical order, other than [Brevibacterium] which is in brackets due to controversy over its taxonomy.

### Effect of Media Selection on Fungal Diversity

Media type affected fungal richness with R2A capturing the most total (Figure 5C) and pyrophilous (Figure 5D) fungal genera whereas MYA captured the fewest. Across all media types from fresh soil, for total fungi we cultured 14 genera (Figure S11), and 29 taxa (Figure S12), and 8 genera (Figure 7B) and 23 taxa for pyrophilous fungi (Figure S13). Among total fungi, R2A media captured the highest richness (57% of genera) while both MYA and PyOM were equally diverse (43% of genera). Among pyrophilous fungi, R2A also captured the highest richness (89% of genera), followed by PyOM (50% of genera), and MYA was the least rich (38% of genera). While PyOM did not capture the most diversity, it was the only media to capture *Neurospora* (Figure 7B), a well-documented post-fire fungus with relatively little knowledge about its ecology (54).

## Discussion

Here, we present results of an experiment to test soil storage and media selection to obtain the best diversity and abundance of bacterial and fungal isolates, which have informed our continual culturing efforts resulting in a collection of 544 pyrophilous microbes (286 bacteria; 258 fungi) (Table S3). To our knowledge, this is the first study examining the impact of soil storage on fungal diversity, and this is also the first large-scale culturing effort of pyrophilous bacteria, and among the largest soil and mushroom based culturing efforts of pyrophilous fungi since 1910 (22). For both bacteria and fungi, freezing soil without glycerol yielded the highest number of pyrophilous genera on rich media, and from fresh soil, R2A yielded the most genera. Although it did not capture the highest number of CFUs or highest number of taxa, novel solid agar media we created from field-harvested PyOM was necessary to capture some key pyrophilous genera of interest like *Neurospora* for fungi and *Noviherbaspirillium* for bacteria.

For bacteria, we conclude that freezing soil at -80°C is the best storage method relative to fresh soil, yielding the highest number of CFUs and total and pyrophilous bacterial genera when plated onto rich media. Interestingly, adding glycerol increased the number of taxa but reduced the number of total and pyrophilous genera isolated. While CFU based analyses are useful in giving an approximation of the total number of culturable cells present in the original sample and a broad overview of the effectiveness of certain culturing practices at cultivating cells, it does not capture the important distinction of isolate diversity. However, in this case, freezing soils without glycerol captured both the highest number of CFUs and yielded the vast majority of the pyrophilous bacterial genera of interest based on the post-fire bacterial succession literature (45–47, 49, 55, 56). While culturing from fresh soil did capture the greatest number of total genera, freezing soil captured the greatest richness of pyrophilous genera (Figure 4). Interestingly, freezing soil resulted in more total and pyrophilous bacterial genera being isolated, but adding glycerol increased the total number of taxa and was necessary to isolate *Massilia,* which is a bacterial genus repeatedly observed post-fire (46, 47, 49, 57), with very little understanding about its ecology, making it a prime candidate for further study. It is possible that taxa only isolated when glycerol was added could be either taxa that are more sensitive to the thermal shift that occurs with freezing or they could be taxa which do not sequester themselves into aggregates and other microenvironments and therefore need the glycerol to buffer them against lysing under freezing conditions. Therefore, it may be of value for researchers to freeze some soil with glycerol as well as without, to capture some of these more potentially vulnerable taxa. Finally, dried soils performed the worst in capturing total and pyrophilous bacterial diversity, which surprised us given how dry post-fire soils usually are. While there is little similar existing literature to compare to, our findings differ from one previous study that found that refrigeration at 4°C was better than freezing at -20°C at retaining microbial biomass and respiration (33). In contrast to our burned chapparal soils, this study examined unburned grassland and forest soils. Different soil structure (prevalence of soil aggregates, water holding capacity, porosity, etc.) can influence microbial activity (58) and therefore could influence how soil microenvironments behave under storage. Although we know of no other cultivation studies to compare to, a study examining how storage time (0-14 days) and temperature (room temp, 4°C, and -20°C) affected 16S sequencing results similarly found that freezing preserved the most bacterial diversity, though they did not test freezing to -80°C as we did (34). Both of these studies only froze at - 20°C rather than -80°C, which may have resulted in fewer taxa retained (59).

Soil storage method seemed to matter less for pyrophilous fungi than bacterial genera. While there was one method that stood out as the worst (freezing at -80°C with glycerol), the remaining methods captured most of the richness observed (Figures 4D and 6B). This could be due to the resilience of fungal spores to a variety of disturbance effects (60), but it is also possible that MYA was not the ideal culture medium for pyrophilous fungi, given the success of R2A and PyOM media at cultivating pyrophilous fungi (Figures 5D and 7B). Future work repeating the storage experiments with R2A and PyOM might improve pyrophilous fungal isolate retention. Despite this caveat, we were able to draw some very surprising conclusions from our experiments. For example, our hypothesis that air-drying soil would be the most effective storage method was fully unsupported by our results. Indeed, air-drying yielded the lowest CFU counts and total and pyrophilous diversity. This is an important finding since drying soils prior to laboratory use is not uncommon (34, 61, 62). Notably, we were unable to find other studies exploring the impact of soil storage on fungal communities, indicating a strong need for these methods to be tested across kingdoms.

Surprisingly, R2A was the most effective culturing media for capturing pyrophilous bacteria and fungi. R2A produced the second highest number of CFUs after the rich medias for bacteria (LB) and fungi (MYA) (Figure 3) and the most diverse pyrophilous bacterial (Figure 5B) and fungal (Figure 5D) isolates. Since R2A was developed to culture bacteria from oligotrophic potable drinking water (63), this suggests that many pyrophilous microbes prefer a more oligotrophic environment. While shifting life-strategies of microbial communities under stress can be extremely varied and nuanced (64), oligotrophic bacteria and fungi may be favored under an increasing stress gradient (such as carbon stress, soil moisture stress, or heat stress) (65), which might be ideal for capturing a wide diversity of microbes thriving in the stressful post-fire environment. However, this is also a surprising result as post-fire environments tend to be high in nitrogen and phosphorus, even if the carbon landscape has radically changed (26, 28). It is therefore possible that these pyrophilous taxa are less affected by the high nitrogen and phosphorus and instead conform more to the previously observed pattern of oligotrophy under a stressful environment. As such, it is possible that availability of certain carbon forms plays a more integral role in post-fire nutrient acquisition and should be studied further. While previous studies exist examining the role of media conditions on microbial culturability (18, 32, 66, 67), these studies often take either a fine-scale approach to tuning a specific culture medium or test only CFU-based metrics rather than diversity or use only rich media. Continued exploration of media conditions is essential to increasing the number of cultured microbial species, and insights gained from our study may help open new paths in culture media development.

Finally, although our home-made PyOM media captured the lowest CFUs of bacteria and fungi, and the lowest bacterial genus richness, it captured the second highest fungal richness and was necessary to isolate key bacterial (i.e. *Noviherbaspirillum soli*) and fungal (i.e. *Neurospora*) pyrophilous taxa of interest. In this case, isolating finicky pyrophilous taxa of interest was improved by the low-density growth we observed in PyOM, which likely resulted from the lower nutrient content similar to R2A media. Rich media like LB, MYA, and potato dextrose agar are often plagued by fast growing ruderal species that can outpace taxa that grow at lower density (68). These taxa are particularly interesting because *Noviherbaspirillum* has been repeatedly observed increasing in abundance post-fire (45–47) and recent genomic work has shown *Noviherbaspirillum* to have a diverse array of PyOM degradation pathways (52). Likewise, *Neurospora* is famously found post-fire and is widely used as a genetics model organism despite very little understanding about its ecological role in the environment (54) though recent genomic work has also characterized potential PyOM degradative abilities (53). Therefore, we conclude that R2A is best media to use to maximize pyrophilous bacterial and fungal diversity, but it would be ideal to also include a PyOM media if specifically interested in isolating the most finicky and interesting pyrophilous microbes.

While we made significant progress regarding testing effective culturing practices for environmental microbes and specifically pyrophilous bacteria and fungi, much additional work remains and caveats exist. Except for the home-made PyOM media, we limited ourselves to commonly available laboratory culturing media. We acknowledge there is a near endless range of possible medias to use (66), possible range of growth temperatures (50, 66), additional isolation strategies (69), and media pH (17, 67), which might be more effective for the culturing of pyrophilous microorganisms. However, we assert that the broad trends (rich vs. oligotrophic vs. PyOM media) yield valuable insights into the culturing of these microorganisms. Additionally, as this work is intended as a foundation for understanding the culturing of pyrophilous microbes, we endeavored to stick to laboratory materials and methodologies that are commonly available to a wide range of research institutions, and accessible for undergraduates, which included most of our co-authors. The PyOM media uses a water-extraction based method of incorporating nutrients, which might have precluded key nutrients required for post-fire metabolism that are less water-soluble in PyOM than others (70), which might have missed some pyrophilous microbial taxa. While we have some limited data on our PyOM media composition (GC mass spectrometer analysis showed 10x more nitrogen present in PyOM compared to LB), additional metabolic analysis is needed to fully characterize this environmentally-derived media. Additionally, the culturing effort in this study all took place aerobically at room temperature in a temperature controlled lab. We also acknowledge that our CFU calculations are not normalized by dry soil weight, since we instead prioritized the preservation of potentially sensitive taxa over drying soil prior to calculation, which may limit direct comparisons from our CFU data to that of other studies. Finally, while we acknowledge that separating microbial cultures based on colony morphology is difficult at best (71–73), we decided that choosing colonies based on morphotype and then identifying with Sanger sequencing was an acceptable concession to the technical and logistic demands of isolating, stocking, and identifying every single colony we grew. This does mean that we could have missed some species-level diversity since some species can share extremely similar colony morphologies (72, 73) and techniques like dilution to extinction culturing might capture more isolates identified to species level (74–76). However, with the difficulties of identifying taxa to species level with rRNA fragments alone coupled with the potential morphological limitation just discussed, we believe our genus-level observations to be more robust hence our exclusion of species-level analyses in this study. Despite these caveats, we still obtained a wide diversity of total (Table S1) and pyrophilous (Table S2) bacteria and fungi and found meaningful differences in media and storage types.

## Conclusions

Here, we present the first study to systematically test how soil storage methods impact fungal cultivability, and the largest scale test of how soil storage and media type affect bacterial and fungal cultivation. We found that storing soils at -80°C without glycerol and utilizing a combination of rich media and oligotrophic R2A media preserved the greatest diversity of culturable pyrophilous bacteria and fungi compared to fresh soils from burned chaparral. However, certain pyrophilous taxa of interest were only captured from soil frozen with glycerol, like *Massilia*, and on our novel PyOM media, like *Neurospora* and *Noviherbaspirillium*, indicating a potential value in using two media and storage types. This experiment contributed to a larger culturing effort resulting in a collection of >540 pyrophilous microbes (286 bacteria; 258 fungi) isolated from soil, smoke, and mushrooms isolated from 7 California wildfires (Table S3), which can be used for future multi-omics, phenotypic, and applied microbiological research. Indeed, we have already begun investigating the traits and trade-offs of these cultures through both phenotypic and genomic profiling (52, 53, 77). Sequencing genomes from isolates with concurrent biophysical assays is a powerful yet under-used tool to gain insight into microbial traits and connect genomic information to trait expression (78). Obtaining the cultures is the first step to characterizing their traits via ‘omics and phenotypic methods.

## Materials & Methods

### Site Description

To test the best soil storage method and media to cultivate pyrophilous microbes, we sampled from The El Dorado wildfire, which burned ∼92km^2^ Manzanita (*Arctostaphylos)* dominated chaparral shrublands, in Yucaipa, CA from September through November of 2020. We examined high-resolution successional data of bacterial and fungal communities after a similar nearby high-severity Manzanita dominated chaparral wildfire (46), and compared to other post-fire studies that had successional data (47, 55), to identify 6 months post-fire as the ideal time to attempt to isolate a wide range of pyrophilous bacteria and fungi. We specifically identified the timepoint as being a point during microbial succession where several hypothesized pyrophilous groups (fast-growers, post-fire nutrient acquisitive microbes, and thermotolerant microbes) (45, 46, 49) were in proportional flux but all present in varying amounts (46). We thus collected samples from high severity burned soils in June 2021, 6 months post-fire. Burned sites were characterized as having 50 to 75 percent slopes with dry, sandy soils in the Springdale family-Lithic Xerorthents association (https://websoilsurvey.nrcs.usda.gov/app/). At the time of collection soil moisture content was 9.72% (±2.7%) per gram soil as assessed by mass loss after drying.

### Soil Collection and Storage

Approximately 250mL of soil from 0-10 cm below the ash layer of burned soil was collected from 9 plots with ethanol cleaned releasable bulb planters and placed into whirl-pak bags 6 months post-fire following methods used in a similar ecosystem (46). After collection, soil was 2mm sieved and either used as fresh inoculum for cultivation or stored in four different ways: air dried, refrigerated (4°C), frozen (-80°C), or suspended 50:50 %w/v soil with glycerol to create a soil slurry and then frozen (-80°C) (Figure 1).

### Culturing Medias Used

Fresh soil samples were cultured immediately onto rich (LB for bacteria, MYA for fungi), oligotrophic (R2A), or PyOM media. After 3 months of storage, stored soil samples were cultured onto rich media. This means that the cultures isolated from fresh soils on rich media served as both the “rich media” condition for the media experiment and the “fresh” comparison for the soil storage experiment (Figure 1).

We produced LB broth by adding 15g of powdered LB Broth (Fisher Scientific, CAS# 73049-73-7) and MYA broth by adding 5g each of powdered Malt Extract (Research Products International, Genessee, CAS# 8002-48-0) and Yeast Extract (Sigma-Aldrich, Cat # 70161) per liter of DI water. For solid agar media, 15g per liter of bacteriological grade agar (Apex Chemicals, Cat # 20-273) was added to either the LB or MYA broth. R2A agar was produced using powdered pre-mixed R2A agar (Criterion, Cat # 16721) following manufacturer directions.

We developed an in-house recipe of PyOM media using biochar harvested from the El Dorado Fire burn sites (Figure S1). Char pieces that were uniformly black in color, brittle, and appeared fully pyrolyzed were selected from the same plots where soil samples were taken. First, we ground biochar into a fine powder with a coffee grinder (Chefman, Model: RJ44-OPP-RED) for 3-5 minutes at maximum speed. Then, we added 50g of ground biochar to a 2L Erlenmeyer flask, added 1200mL of DI water, covered the mouth of the flask with a watchglass, stirred on a hotplate for 24 hours at 40°C, then removed from heat to settle for 8 hours. We then siphoned liquid contents into a clean Erlenmeyer flask, being careful to not disturb the bottom sediment. We passed the liquid through cheesecloth, and then twice through a coffee filter to remove any remaining sediment before transferring to clean containers for autoclaving. We added 13g of bacteriological grade agar (Apex Chemicals, Cat # 20-273) to each liter of PyOM media, since adding 15g resulted in under-dissolved agar in the final product. This resulted in a media that should reflect the more complicated carbon sources found in char (PAHs, benzoic compounds, etc.) and contained almost 10x more nitrate than LB as assessed by GC mass spectroscopy (LB avg NO_3_^-^ conc. = 0.27mg/L, PyOM avg NO_3_^-^ conc. = 1.2mg/L). We used environmentally sourced char from the El Dorado burn site, which we believed was the best representative substrate for pyrophilous microbes, but since there can be fluctuations in chemical content across char collections, the PYOM media used in this study was derived from a single acquisition of char. We used DI water for all media types (LB, MYA, R2A, PyOM) and all were autoclaved at 121°C for 45 minutes.

We added antimicrobials to the liquid agar after autoclaving, once it had cooled to 60°C. For media used to isolate fungi and intended to resist bacterial growth, we added filter sterilized gentamycin and ampicillin (Hardy Diagnostics, Santa Maria, CA) to a final concentration of 10mg/mL each. For media used to isolate bacteria and intended to resist fungal growth, we added filter sterilized cycloheximide (Research Products International, CAS # 66-81-9) to a final concentration of 20mg/L.

### Experimental Design and Microbial Isolation

We collected 12 soil samples from high-severity burn locations (3 plots with four 1 m² subplots each). Three grams from each sample were combined into a single homogenized soil sample, which was then subsampled for culturing treatments: fresh soil plated on LB, MYA, R2A, or PyOM media, or soil stored for three months under different conditions (air-dried, refrigerated at 4 °C, frozen at −80 °C, or frozen at −80 °C with glycerol). For samples stored in glycerol, 1g of fresh soil was mixed with 1mL of glycerol resulting in an approximately 1:1 w/v soil to glycerol slurry prior to freezing. Stored soil was only plated onto rich media to not introduce additional confounding variables of media type into the soil storage experiments. For each treatment, 1 g of soil was suspended in 5 mL sterile DI water in four replicate Falcon tubes, serially diluted (1:10–1:10,000), and 100 µL of each dilution was plated onto 10 replicate plates targeting either bacteria or fungi. All fungal plates contained ampicillin and gentamycin, while bacterial plates contained cycloheximide. In total, 560 plates were prepared (2 kingdoms × 4 dilutions × 10 replicates × 7 treatments plus sterile controls). Plates were incubated at room temperature (∼25°C) for ∼1 week, after which colony-forming units (CFUs) and unique morphologies were counted. Plates containing 30–500 colonies per plate were considered countable, yielding 6–9 replicate plates per treatment. Unique bacterial and fungal morphotypes were isolated, DNA extracted, and Sanger sequenced, and isolates stocked (Figure 1). Diversity was assessed using genus richness (a measurement of α-diversity) of total or pyrophilous genera captured by each treatment.

Microbes were morphotyped by size, color, and growth rate. For bacteria and yeasts, individual colonies were picked with sterile pipet tips and streaked onto fresh LB (without antifungals). For filamentous fungi, a 1 cm² agar square from the colony edge was transferred onto fresh MYA (without antibacterials). When microbes failed to establish on LB or MYA, they were maintained on their original media (R2A or PyOM). Isolates were incubated at room temperature for 1 week and re-isolated until three consecutive clean cultures were obtained for DNA extraction and stocking.

For long-term storage, bacterial isolates were grown in LB broth (3 mL) at 180 rpm for 24–72 hrs until turbid, then mixed 1:1 with 40% sterile glycerol (final glycerol concentration of 20%) in cryovials and stored at −80 °C. Fungal isolates were grown to within 1 cm of the plate edge, then 5 mm agar plugs from the colony margin were transferred into screw-cap tubes containing 1 mL autoclaved tap water (79). Duplicate stock tubes were prepared for each isolate.

### DNA Extraction

DNA was extracted from all unique isolates using home-made Extract-N-Amp Extraction Solution (80) made by combining 10 ml of 1M pH 8.0 Tris-HCl, 1.86g KCl, 0.37g EDTA, and 80 mL DI water, then adding 1M NaOH until pH 9.5 and then diluting to a total volume of 100mL and filter sterilizing. For bacteria, one well-isolated colony was picked using a sterile 10µL pipet tip, and for fungi, a 1mm^2^ area of the growing edge of the fungus was scraped with a flame-sterilized probe. The tip or probe was then submerged and swirled in 10µL of Extraction solution, which was heated at 65°C for 10 minutes followed by 95°C for 10 minutes in a Thermocycler (BioRad). Once finished, 10µL of 3% BSA was added and the samples were left to chill overnight at 4°C before being PCR amplified or frozen at -20°C.

### PCR and Sanger Sequencing

The bacterial 16S rRNA region was amplified using a PA forward and PH reverse primer (Edwards et al., 1989) and the fungal ITS region was amplified with the ITS1F forward (Gardes & Bruns, 1993) and ITS4 reverse primer (White et al., 1990). Each 20 µl PCR reaction contained 13.64 µl of molecular grade water, 0.16μL of Taq polymerase (New England Biolabs, Ipswitch MA) and 2μL of 10x Taq Polymerase Buffer containing MgCl_2,_ 2μL of 10x solution of equally mixed dNTPs, 0.6μL of each primer at 10μM, and 1μL of DNA. Thermocycler conditions were as follows: 94℃ for 2 minutes, then 35 cycles of 57℃ (for fungi) or 62℃ (for bacteria) for 1 minute, 72℃ for 1 minute, then 94℃ for 1 minute, ending with a final extension of 72℃ for 8 minutes (81). The DNA was cleaned using 3μL of diluted Exosap mastermix (Applied Biosystems, Waltham MA) per 7μL PCR product before diluting and preparing for Sanger sequencing. Samples were Sanger sequenced in the forward direction at the UCR Institute for Integrative Genome Biology Genomics Core services.

### BLAST Identification

Raw sequences as .ab1 chromatogram files were trimmed in either SnapGene or Geneious Prime using the same parameters to remove low quality regions (Phred scores less than 30, preserving the largest continuous region of good quality sequence) then saved as FASTA files. Taxa were identified using NCBI BLASTn by generating hit tables of the top 10 matches, with IDs assigned based on query cover, e-value, percent identity, and consensus among hits of similar quality. Isolates were identified to species if query cover was 100%, e-value < 0.01, and percent identity >98%; to genus at ∼95%, family at 90%, class at 85–90%, and phylum at <85%. However, given the difficulty of identifying species with only rRNA fragments, genus level identification is the most reliable and species identifications reflect our best estimation of taxon identity. All isolate names were preserved from the best-matching BLAST identification, including any additional modifiers indicating potentially problematic taxonomy.

### Statistical Analysis

All statistical analyses and figures were produced in R 4.0.2 (82) and all scripts are available on GitHub (https://github.com/DylanEnright93/Culturing_Pyrophilous_Microbes). Normality was tested with a Shapiro test, then a two-way ANOVA was used to test the effect of soil storage or media treatment on bacterial and fungal CFU counts, followed by a post-hoc Tukey HSD test. Heatmap-like visualization of genera and species level diversity were created using the “geom_tile” function in ggplot2 (83). Diversity among treatments was evaluated using α-diversity by calculating the richness of genera per treatment and comparing the percent of total isolates or total pyrophilous isolates (based on literature support) obtained per treatment.

### Data Availability

All bacterial 16S and fungal ITS FASTA sequences are accessioned under accession numbers PV662515 - PV662757 and PV620023 – PV620259, respectively and will be available at the National Center for Biotechnology at Genbank database at time of publication. All scripts used to analyze the data are likewise available on GitHub at https://github.com/DylanEnright93/Culturing_Pyrophilous_Microbes.

## Acknowledgements

We thank the San Bernardino National Forest for allowing us to collect the soils needed for this study following the El Dorado wildfire. We thank many individuals who contributed to the Glassman lab pyrophilous microbial culture including lab managers Judy Chung, James Randolph, and Maria Ordonez, UCR PhD students Fabiola Pulido-Chavez, Arik Joukhajian, and Mark Yacoub, and UCR undergraduates Justin Diab, Wine-Jie Lipardo, Audrey Reichard, Priscilla Shultz, Jorge Ponce, and Nathan Heger. This project was funded by the Department of Energy BER Award #DE-SC0023127 and United States Department of Agriculture-NIFA Award #2022-67014-36675 to SIG and the NSF GRFP to DJE. We also thank Dylan Enright’s Dissertation Committee members Jason Stajich and Marko Spasojevic and anonymous reviewers for their helpful feedback on the manuscript.

## Competing Interests

The authors declare no conflicts of interest or competing financial interests in relation to the work described.

## Notes

### Competing Interest Statement

The authors have declared no competing interest.

### Summary of Updates

Multiple revisions throughout as requested during peer review. Removal of species-level analysis from main paper. Revision of figures 1, 4, and 5.

