## Supplemental Materials for "Evaluating Best Practices for Isolating Pyrophilous Bacteria and Fungi from Burned Soil"

Figure S1

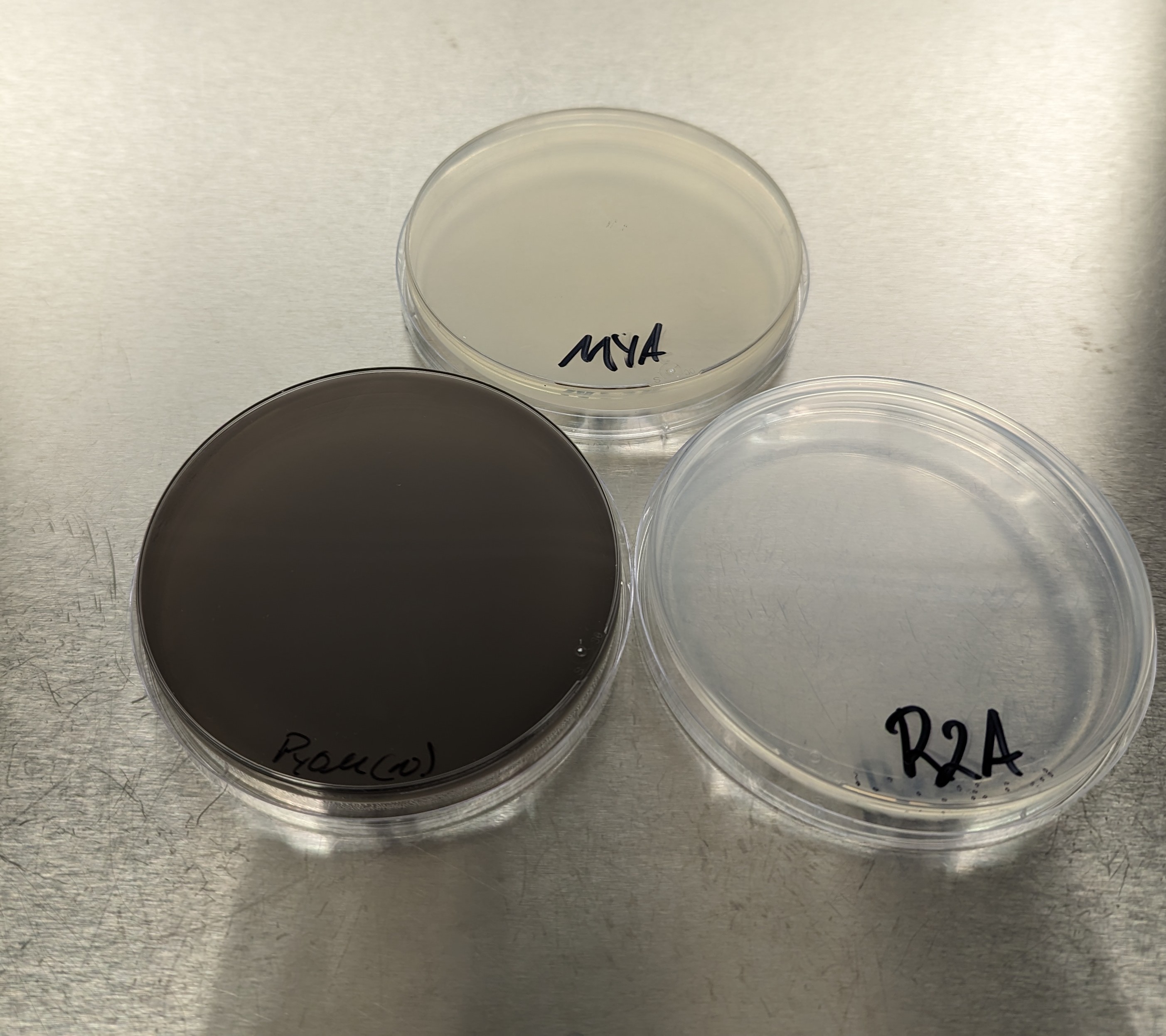

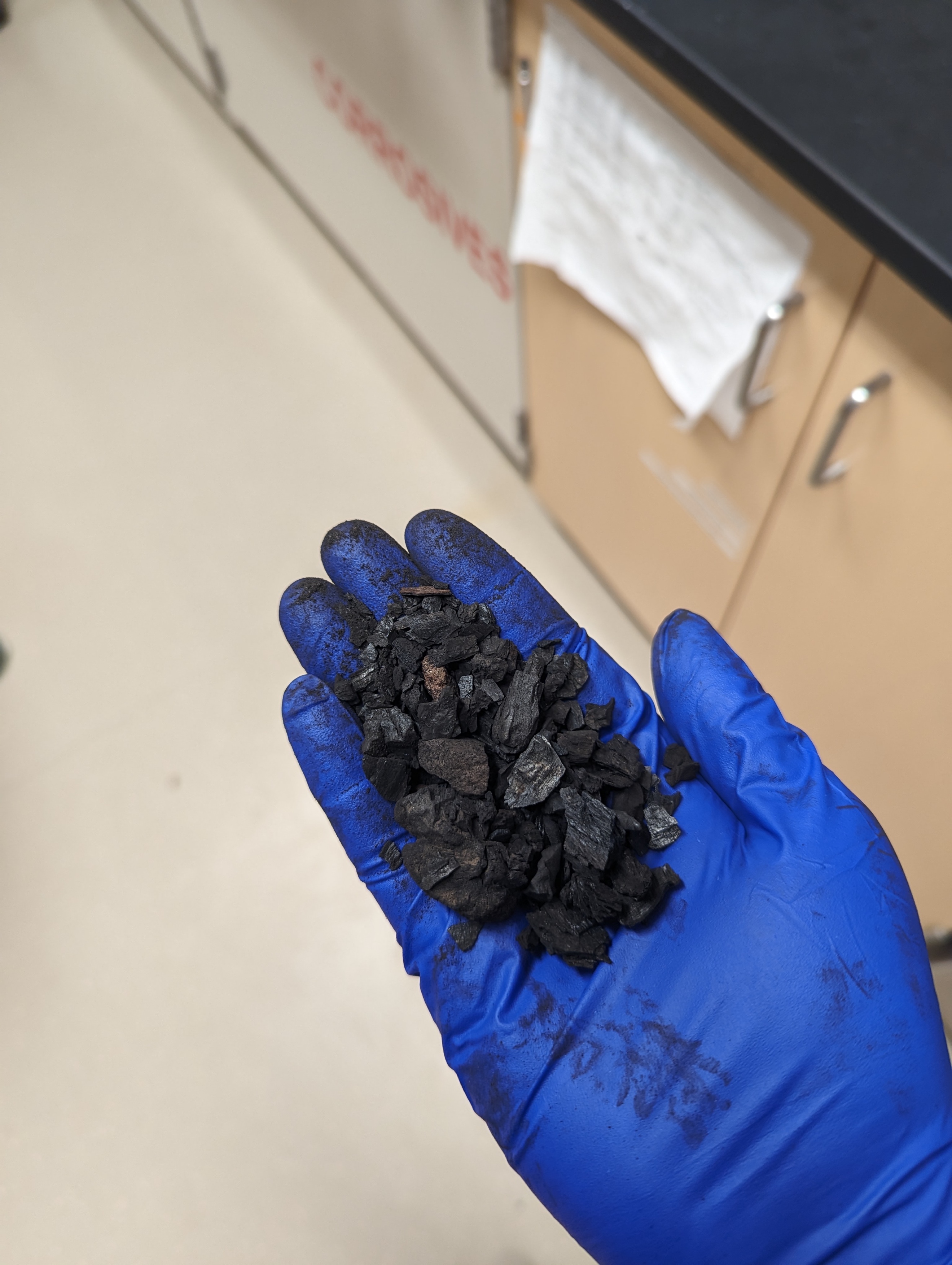

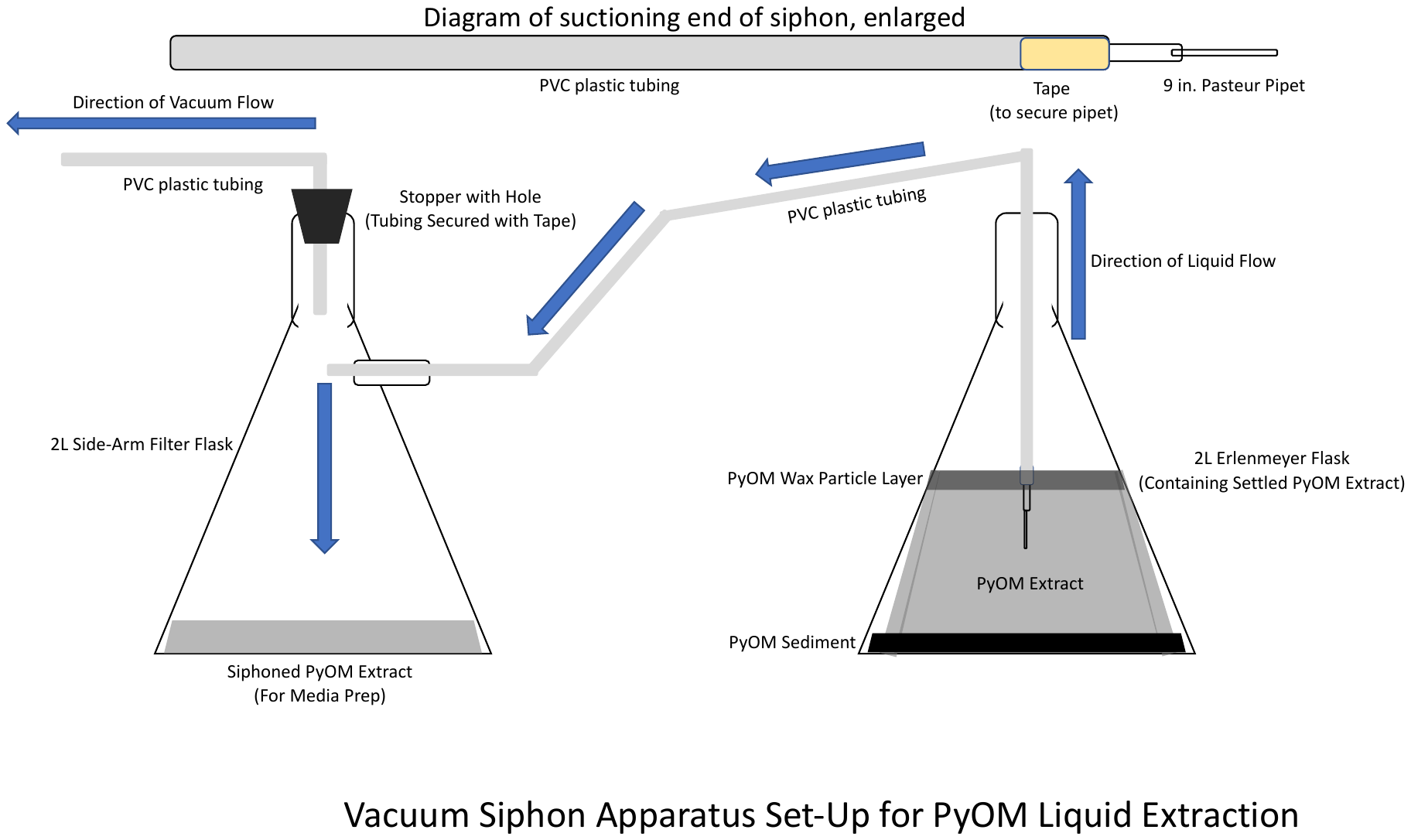

**C**

**B**

**A**

Figure S1: A) Schematic diagram visualizing the vacuum siphoning set up for the creation of PyOM media. Prior to siphoning pyrogenic matter was ground, added to water, and heated at 40ºC overnight. After allowing 1 day of settling, the solution was vacuum siphoned as above. This extract would then be filtered through cheesecloth and coffee filters and allowed to settle one additional time before sterilization. B) And example of the field-sourced pyrogenic organic matter used to make PyOM media. C) An example plate of solid PyOM agar media.

Table S1: All isolates obtained across both experiments.

For supplementary table S1, please see the Microsoft Excel spreadsheet titled “Table S1.xlsx”.

Table S1 shows all isolates obtained from the El Dorado Fire 6 months post-fire across all experimental treatments.

Table S2: All pyrophilous isolates obtained across both experiments.

For supplementary table S2, please see the Microsoft Excel spreadsheet titled “Table S2.xlsx”.

Table S1 shows all pyrophilous isolates obtained from the El Dorado Fire 6 months post-fire across all experimental treatments.

Table S3: All isolates present in the Glassman Lab post-fire microbial culture collection.

For supplementary table S3, please see the Microsoft Excel spreadsheet titled “Table S3.xlsx”.

Table S3 shows all isolates obtained from all fire-related culturing efforts along with information on the substrate the isolates were cultured from and the relative fire information.

Figure S2:

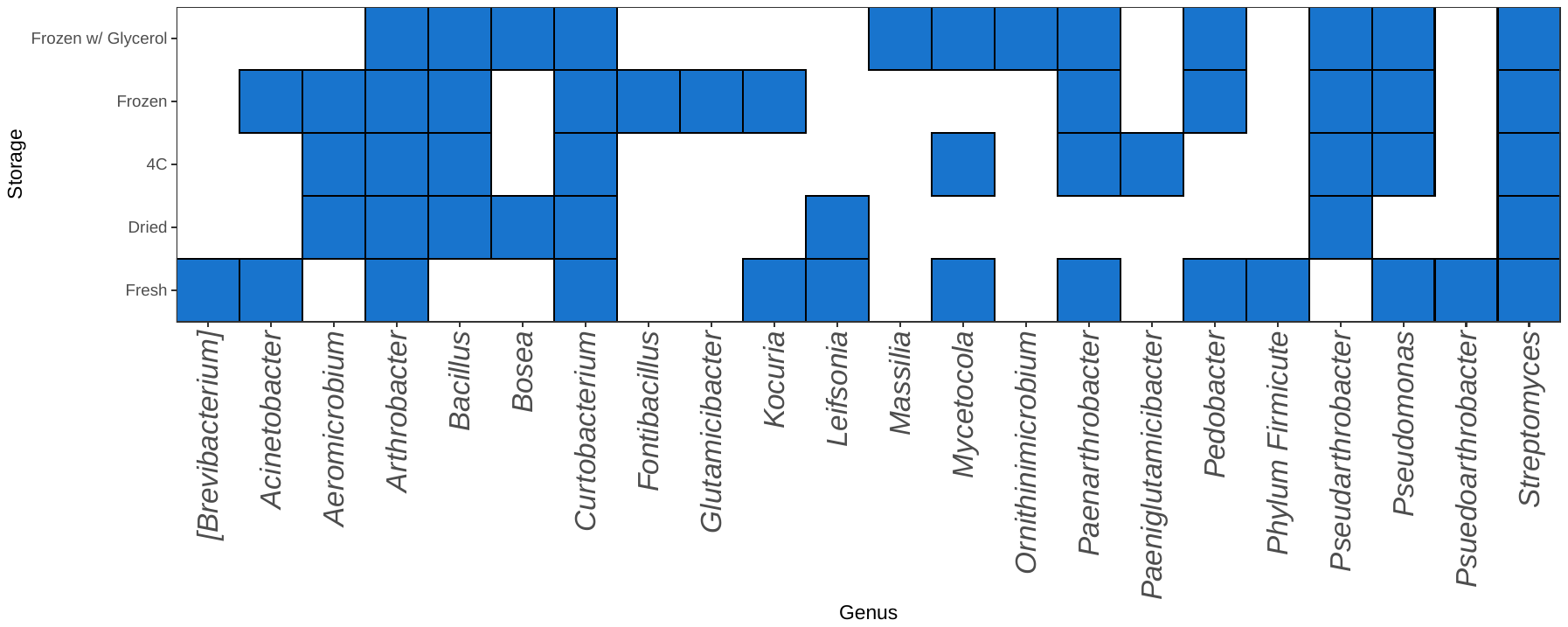

Figure S2: Heatmap-like visualization of the diversity of all bacterial genera across all storage method treatments. Filled blue squares indicate the presence of the genus on the x axis in the treatment on the y axis.

Figure S3

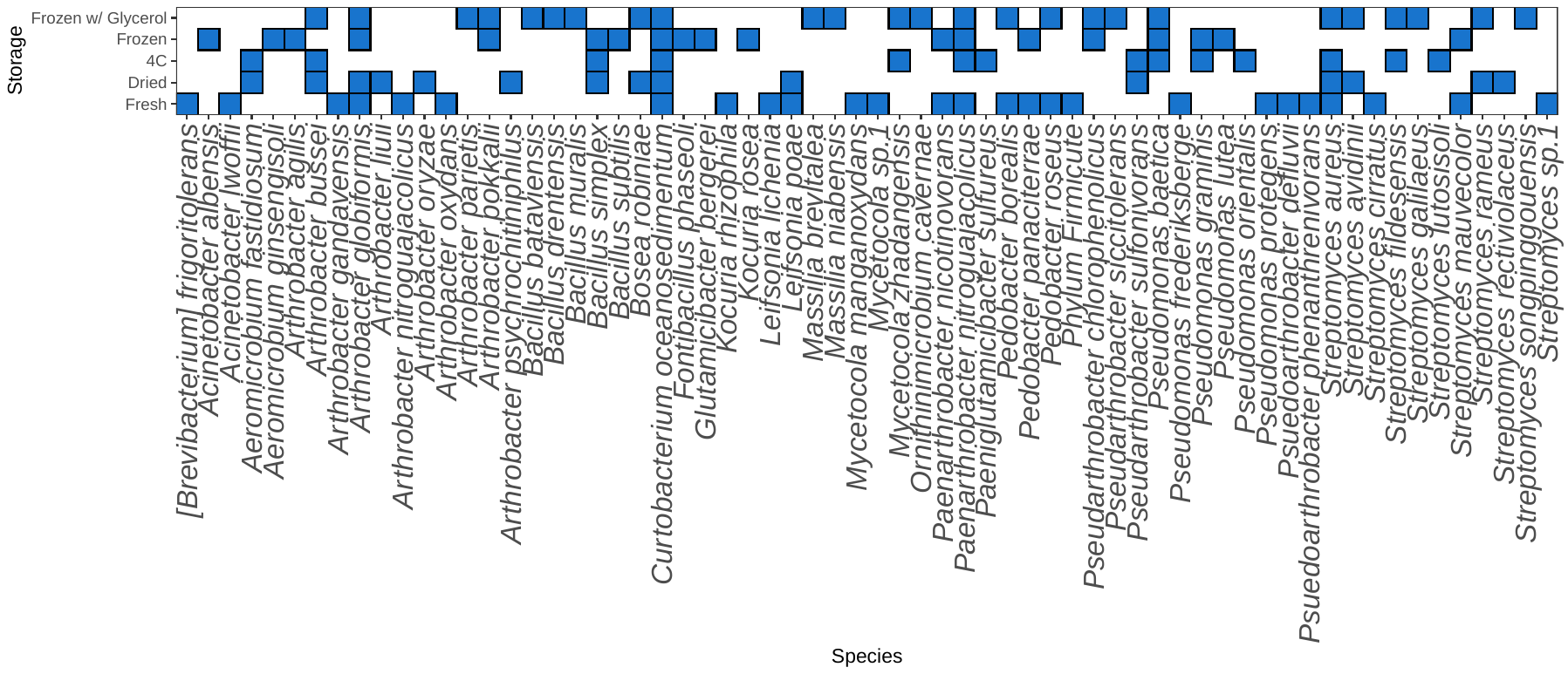

Figure S3: Heatmap-like visualization of the diversity of all bacterial taxa across all storage method treatments. Filled blue squares indicate the presence of the taxa at best identification level on the x axis in the treatment on the y axis.

Figure S4

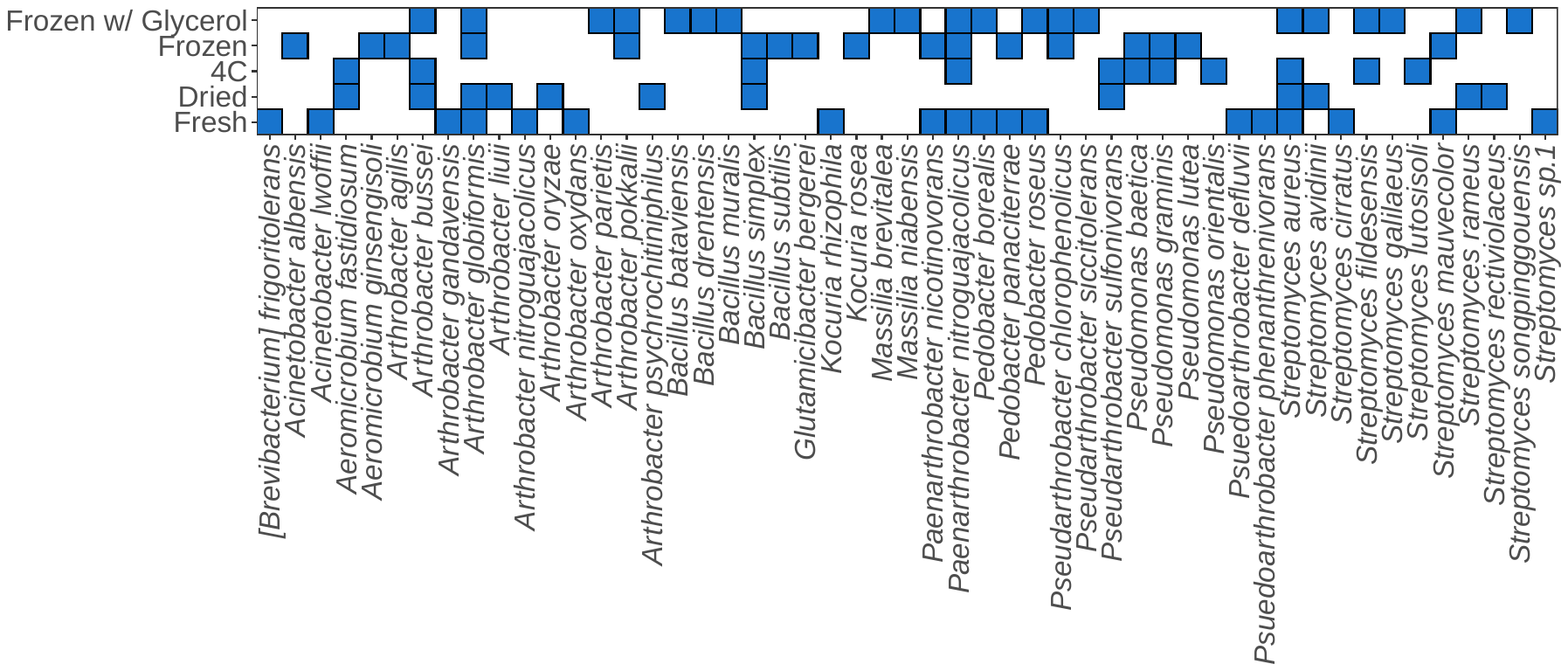

Figure S4: Heatmap-like visualization of the diversity of all pyrophilous bacterial taxa across all storage method treatments. Filled blue squares indicate the presence of the taxa at best identification level on the x axis in the treatment on the y axis.

Figure S5

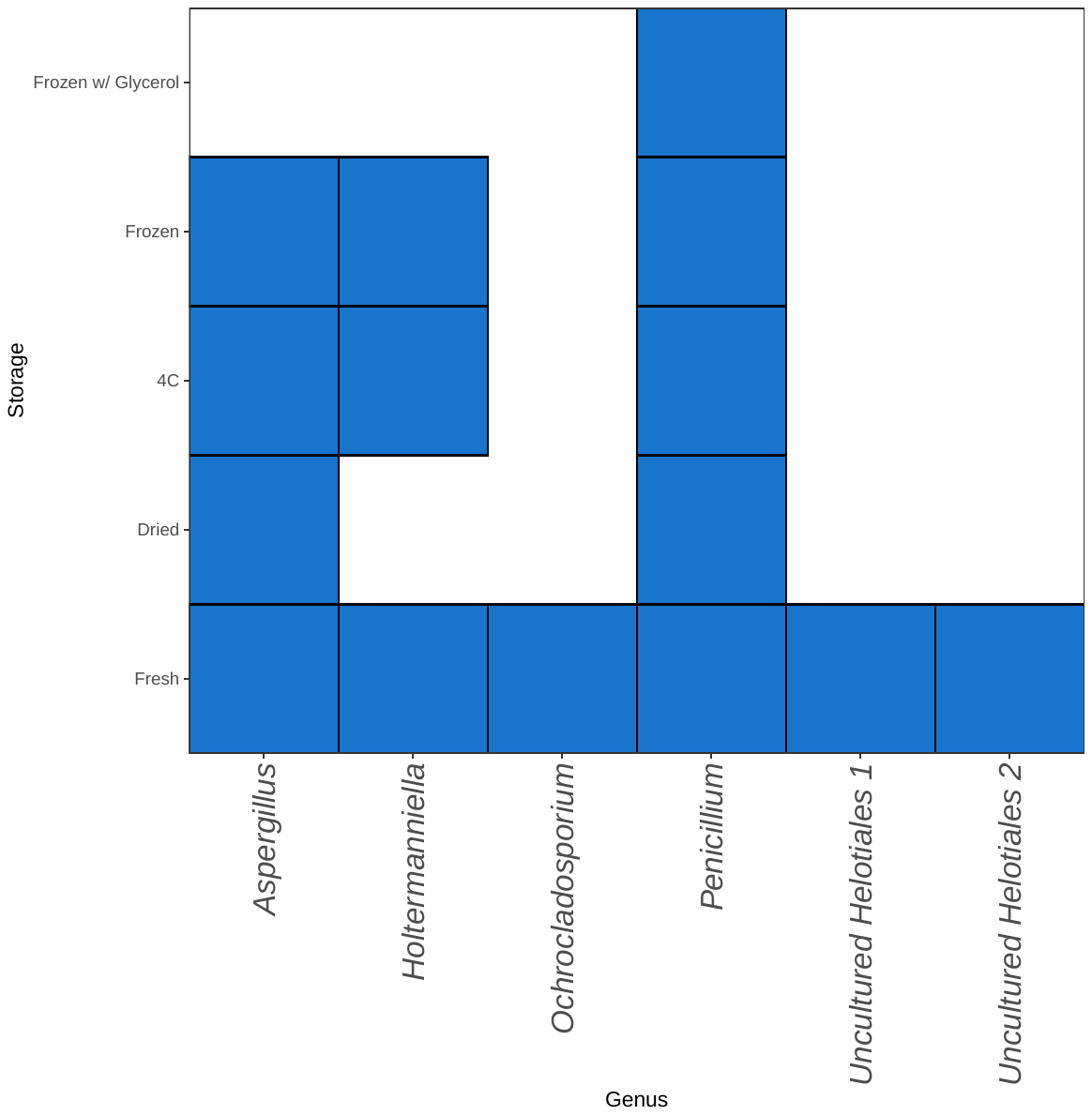

Figure S5: Heatmap-like visualization of the diversity of all fungal genera across all storage method treatments. Filled blue squares indicate the presence of the genus on the x axis in the treatment on the y axis.

Figure S6

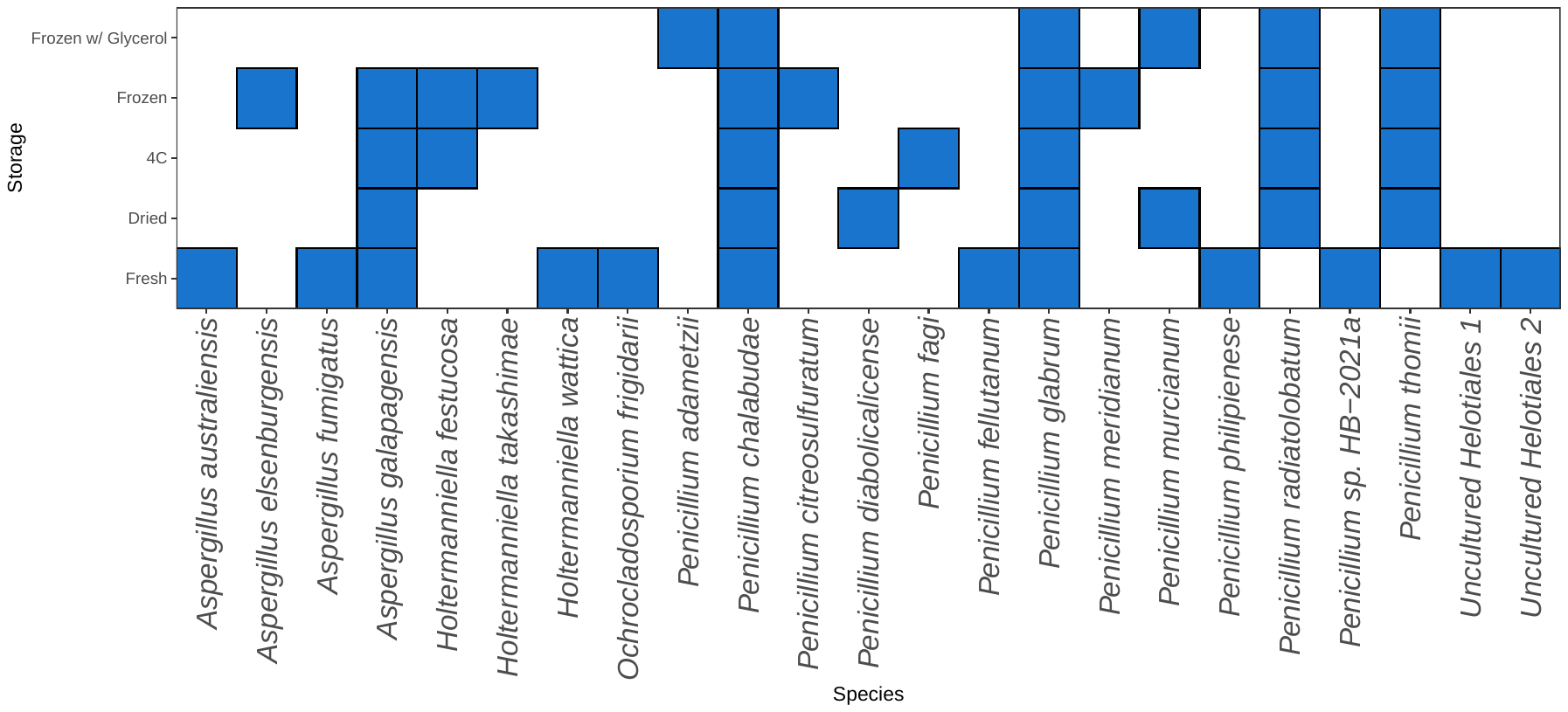

Figure S6: Heatmap-like visualization of the diversity of all fungal taxa across all storage method treatments. Filled blue squares indicate the presence of the taxa at their best level of identification on the x axis in the treatment on the y axis.

Figure S7

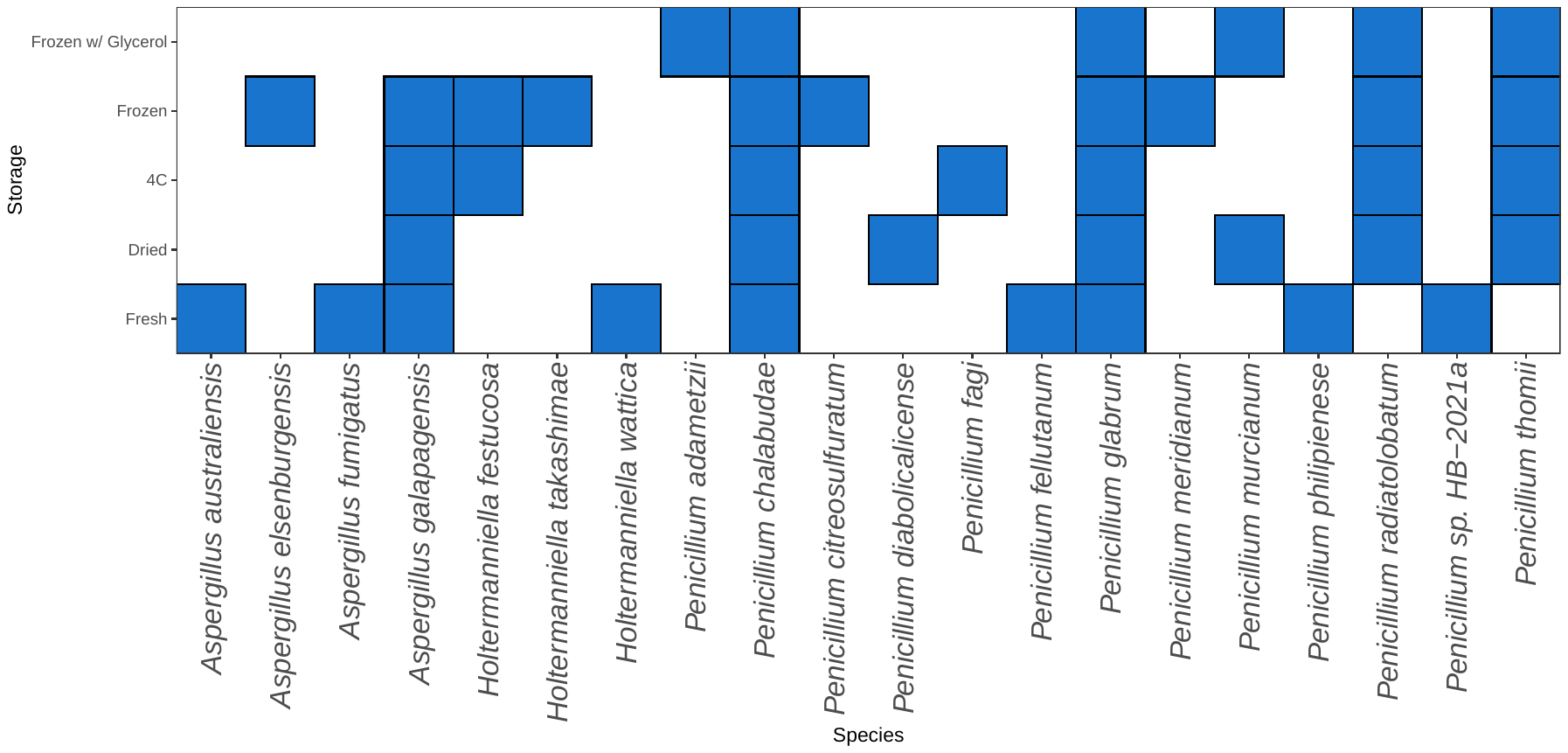

Figure S7: Heatmap-like visualization of the diversity of all pyrophilous fungal taxa across all storage method treatments. Filled blue squares indicate the presence of the taxa at best level of identification on the x axis in the treatment on the y axis.

Figure S8

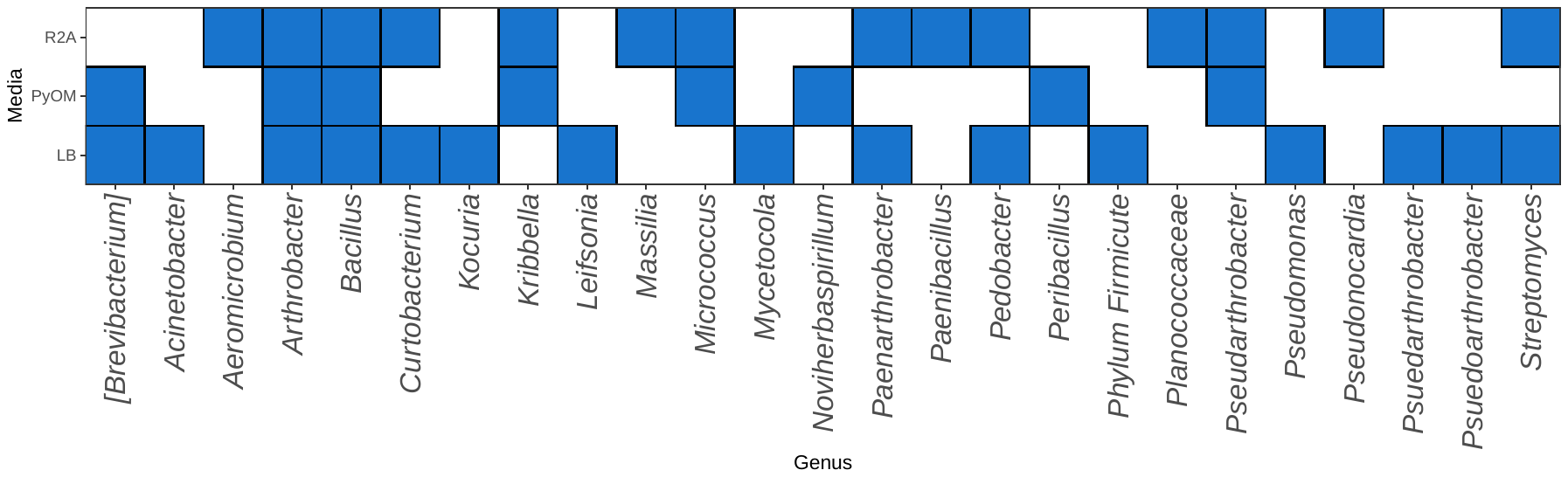

Figure S8: Heatmap-like visualization of the diversity of all bacterial genera across all media type treatments. Filled blue squares indicate the presence of the genus on the x axis in the treatment on the y axis.

Figure S9

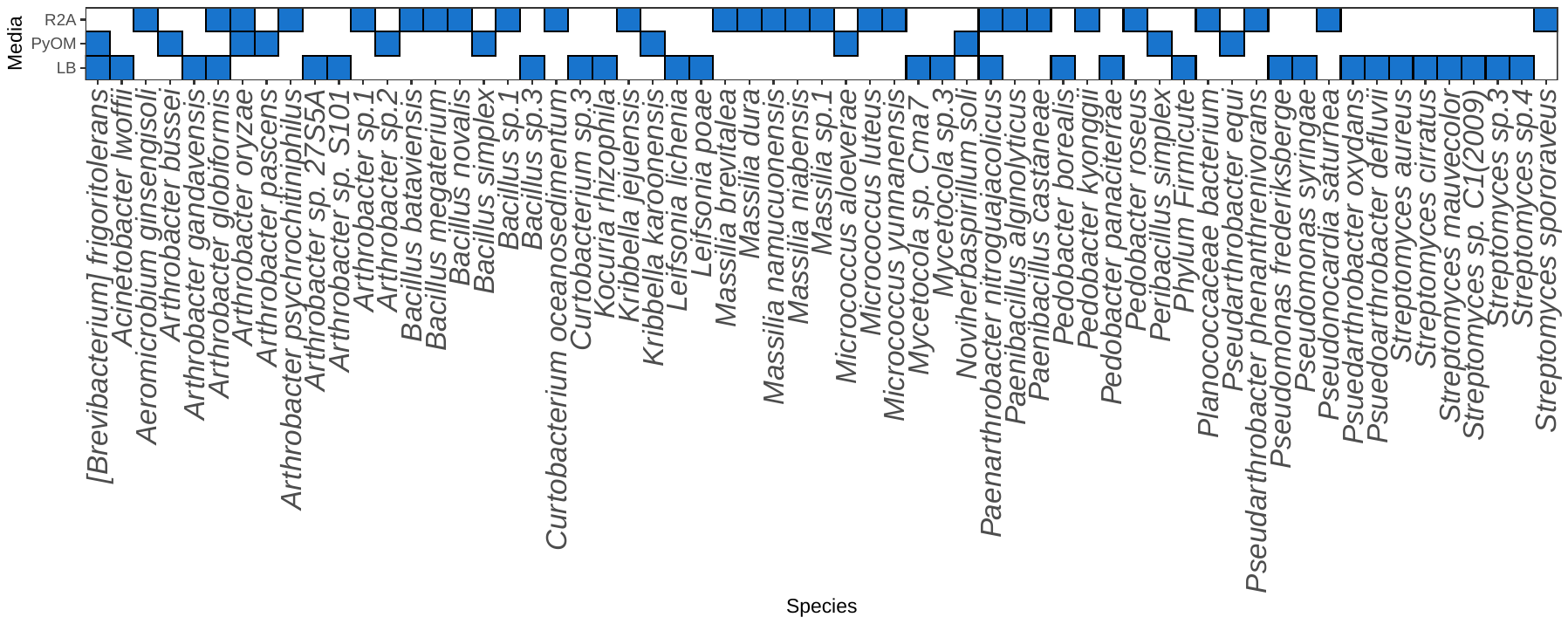

Figure S9: Heatmap-like visualization of the diversity of all bacterial taxa across all media type treatments. Filled blue squares indicate the presence of the taxa at the best level of identification on the x axis in the treatment on the y axis.

Figure S10

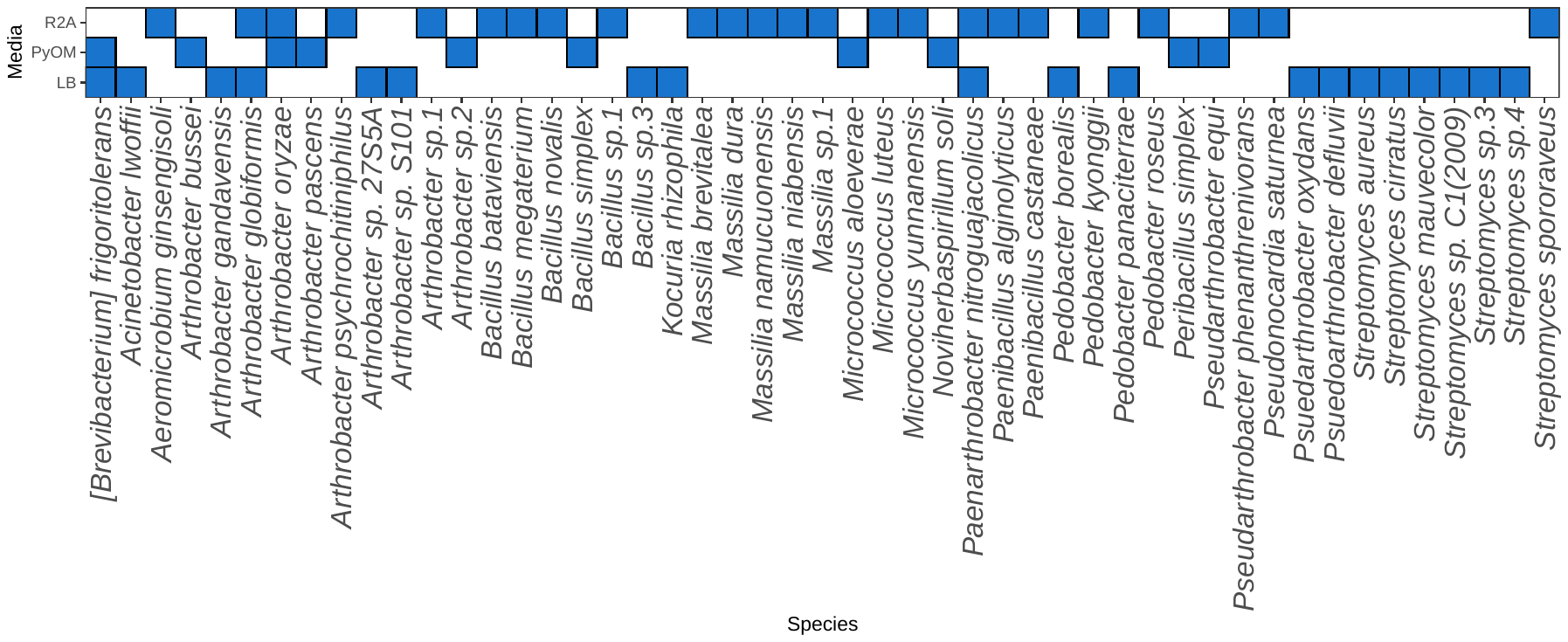

Figure S10: Heatmap-like visualization of the diversity of all pyrophilous bacterial taxa across all media type treatments. Filled blue squares indicate the presence of the taxa at the best level of identification on the x axis in the treatment on the y axis.

Figure S11

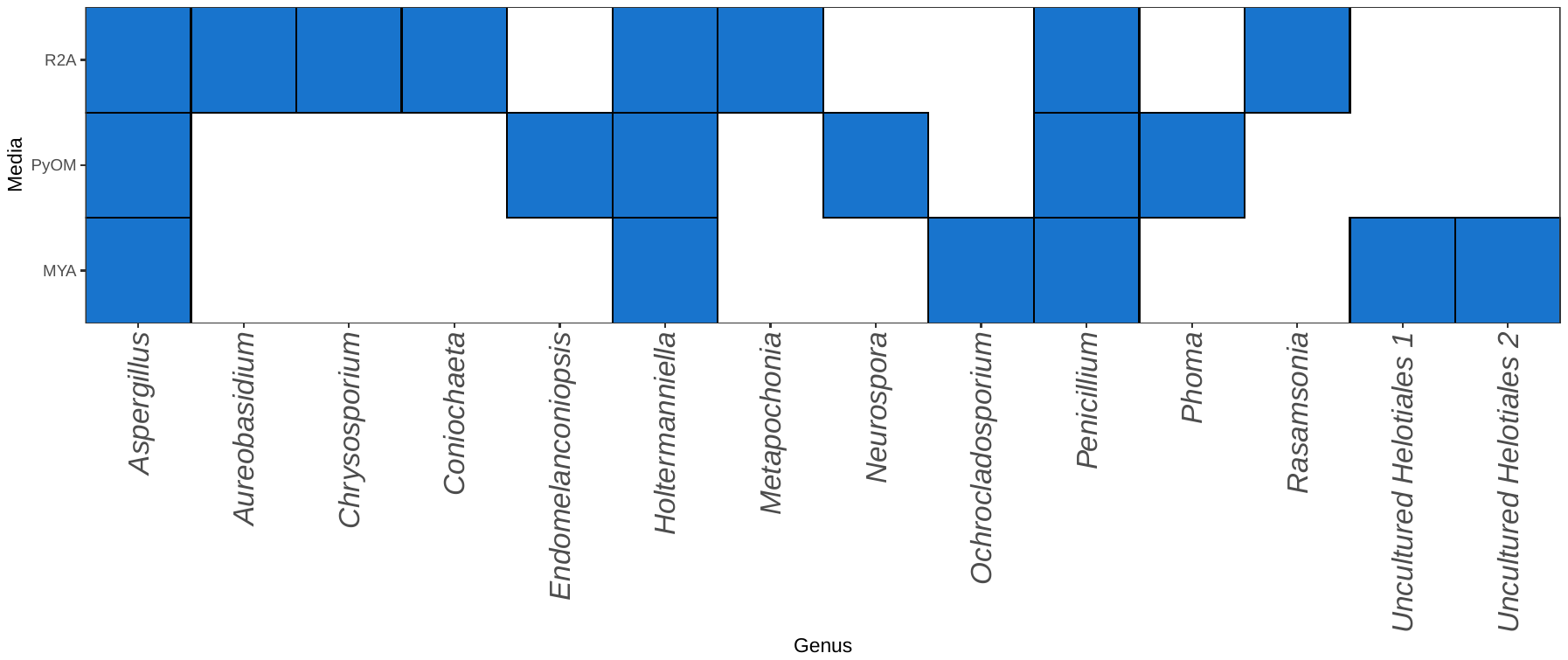

Figure S11: Heatmap-like visualization of the diversity of all fungal genera across all media type treatments. Filled blue squares indicate the presence of the genus on the x axis in the treatment on the y axis.

Figure S12

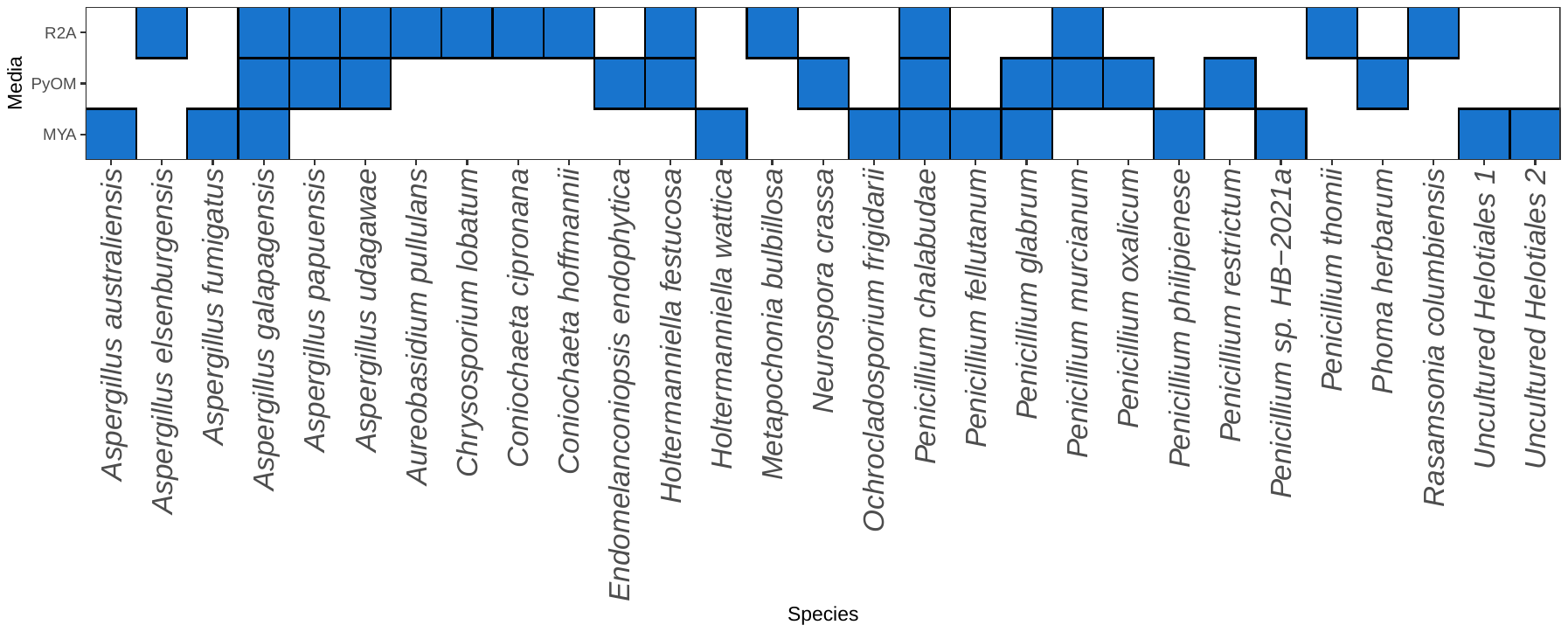

Figure S12: Heatmap-like visualization of the diversity of all fungal taxa across all media type treatments. Filled blue squares indicate the presence of the taxa at the best level of identification on the x axis in the treatment on the y axis.

Figure S13

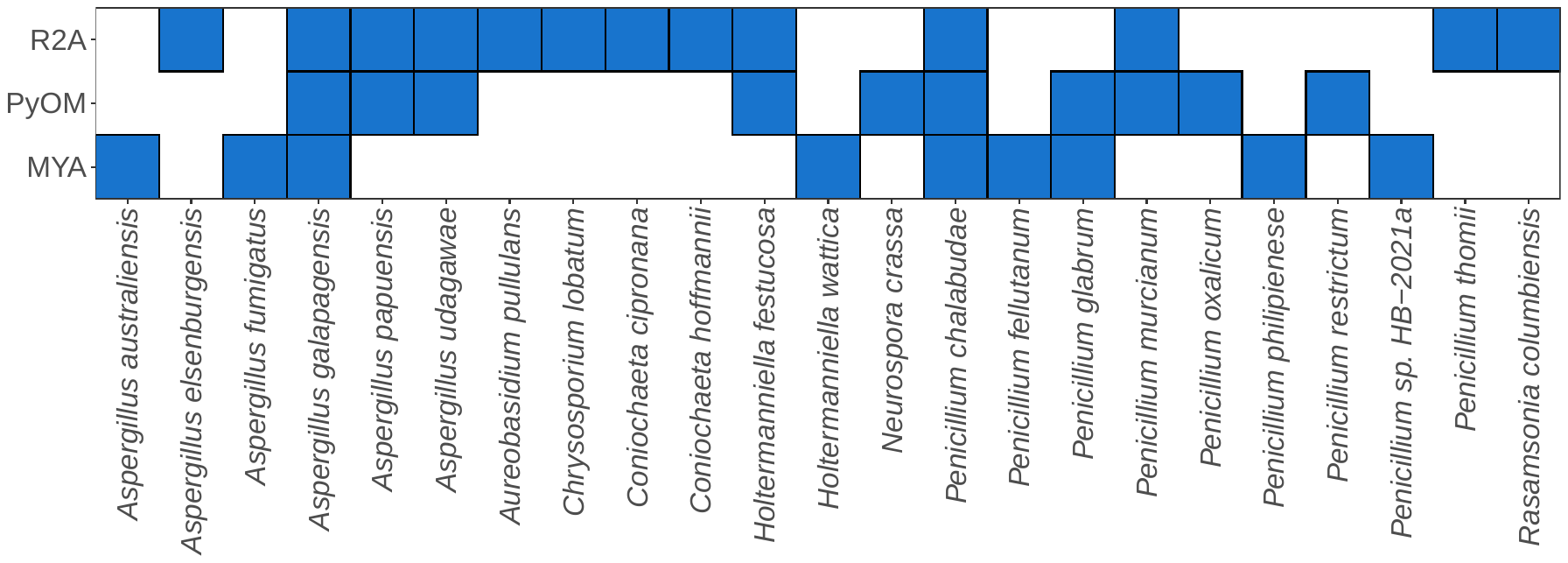

Figure S13: Heatmap-like visualization of the diversity of all pyrophilous fungal taxa across all media type treatments. Filled blue squares indicate the presence of the taxa at best level of identification on the x axis in the treatment on the y axis.
